# Structural dynamics between Argonaute-2 and CK1α promote target RNA release in microRNA-mediated silencing

**DOI:** 10.64898/2026.03.23.713669

**Authors:** Ankur Garg, Leah Braviner, Armend Axhemi, Brianna Bibel, Leemor Joshua-Tor

## Abstract

Argonaute (Ago) proteins associate with 20-22 nucleotide (nt) long microRNAs (miRNAs) to constitute the functional RISC core and downregulate mRNAs containing complementarity to the seed sequence^1–3^. Target RNA engagement in RISC stimulates CK1α-mediated phosphorylation of the conserved eukaryotic insertion (EI) in Ago, releasing the target and enabling the RISC complex to suppress additional target sites for efficient miRNA-mediated silencing^4–6^. Here, we provide a complete structural view of miRNA guide and target binding to human Ago2, showing Ago2 holding the double-stranded guide-target RNA in an untwisted conformation at its center. We visualize the dynamic changes that RISC undergoes as the guide supplementary region progressively base pairs with the target, enabling CK1α binding. Following seed-helix assembly, initial supplementary pairing restricts RISC to a “closed” form, while with half-supplementary pairing, the PAZ domain moves to open RISC to become receptive to CK1α, exhibiting an initial increase in Ago2 phosphorylation. Complete supplementary pairing supports a full PAZ-CK1α interface, allowing for hierarchical phosphorylation of the EI. The combination of target repulsion by EI phosphorylation with an unwound guide-target enables efficient target release to promote RISC turnover.

## Main

RNA interference (RNAi) is essential for development and homeostasis and works by producing a ∼22-nucleotide (nt) long guide microRNA (miRNA) after sequential processing of pri-miRNA hairpins by Drosha and Dicer RNaseIII enzymes^3,7,8^. The guide miRNA is loaded into an Argonaute (Ago) protein to assemble a functional RNA-induced silencing complex (RISC)^9^. Members of the Ago superfamily typically have four conserved structural domains: N, PAZ, MID, and PIWI, with two linker regions (L1 and L2)^10^. The 22-nt length guide is divided into a seed sequence (g2-8), a central (g9-11), supplementary (g12-16), and tail (g17-22) regions. The 5’ and 3’ ends of the guide (g1 and g22) directly bind to the MID-PIWI interface^11,12^ and PAZ domains^13,14^, respectively, allowing the exposed subseed (g2-5) to scan the target mRNA^15–18^, followed by structural changes in Ago that facilitate full complementary seed pairing^19^. Seed pairing further triggers structural changes, promoting supplementary sequence pairing^20^, which allows RISC to recruit accessory factors necessary for effective RNAi^21,22^.

Due to high-affinity binding coupled with a very slow off-rate of miRNA to Ago^23^, RISC can repress countless matching mRNAs, as long as the target mRNA is effectively released post-silencing. This might be challenging when the target RNA, predicted to wrap around the guide twice, isn’t cleaved. Ago proteins are known to undergo different types of post-translational modifications^24^, but a cycle of phosphorylation of the eukaryotic Ago-specific insertion (eukaryotic insertion; EI) by CK1α kinase and dephosphorylation by PP6/ANKRD52 has been attributed to the fast target mRNA turnover by Ago when no slicing occurs^4^. Specifically, in human (Hs) Ago2, the EI (amino acid 820-837) contains highly conserved phosphorylation sites (S824, S828, S831, and S834)^5^. Increased phosphorylation causes a decrease in target RNA association with RISC in cells, and vice versa^4^. A systematic *in vitro* biochemical analysis further revealed that serine residues in HsAgo2-EI undergo CK1α-mediated hierarchical phosphorylation starting with S828 and S831, which further prime the phosphorylation of other nearby serine residues in the EI. Moreover, the study showed that pairing in the supplementary region beyond the g13 position significantly enhanced CK1α-mediated Ago2-EI phosphorylation, which increased the target release rate several fold, thereby revealing a role for supplementary pairing in the target release mechanism^6^. Despite these findings, molecular insights into this crucial step in effective RNAi were lacking.

In this study, we determined several cryo-EM structures of the HsAgo2-miR200b RISC with ZT1 target mRNAs of varying lengths (13nt, 14nt, and 30nt) in complex with CK1α. We found that sufficient base pairing is crucial for HsAgo2 to be in a proper conformation for CK1α docking. In addition, the central region of the guide-target duplex is held by HsAgo2 such that it is unwound. The combination of repulsion of the target by phosphorylation of the EI with the unwound conformation of the guide-target at the center provides a mechanistic understanding of how the mRNA target is released efficiently to recycle RISC and promote further miRNA-mediated silencing.

## Results

### Structural basis of CK1α-mediated RISC^ZT30^ phosphorylation

To investigate the mechanism of CK1α-mediated Ago phosphorylation, we determined a cryo-EM structure of HsAgo2-miR-200b (RISC) with ZT30 (RISC^ZT30^), a 30-nt target corresponding to a natural, highly conserved miR200-binding site in the 3’ UTR of the transcription factor ZEB1, in complex with the kinase CK1α **(Fig 1A-C)**. The RISC^ZT30^-CK1α complex was assembled by combining purified RNA-free^15^ dephosphorylated HsAgo2^6^ **(Supplementary Fig 1A)**, guide, target, and purified CK1α **(Supplementary Fig 1B)** together with AMPPNP. A ∼2.8 Å resolution cryo-EM map clearly shows HsAgo2, loaded with the guide/target duplex (**Fig 1C, Supplementary Fig 1C**). In addition, the portion of the map corresponding to CK1α exhibited high-resolution features for the C-lobe (between 2.8 to 3.5 Å), while the resolution of the N-lobe ranges between 3.5 to 5 Å (**Supplementary Fig 1D**). A significant number of RISC^ZT30^ particles were CK1α-free (**Supplementary Fig 1C**), resulting in a ∼2.8 Å cryo-EM map for the RISC^ZT30^ structure (**Supplementary Fig 1E-F**). We used the crystal structure of HsAgo2/miR122-target complex^20^ (PDB 6N4O) without the RNA, and the crystal structure of CK1α ^25^ (PDB 6GZD) for initial model building, while the guide and target RNA were built de novo into the cryo-EM map. The final model statistics are summarized in **Supplementary Table 1**.

**Figure-1.**
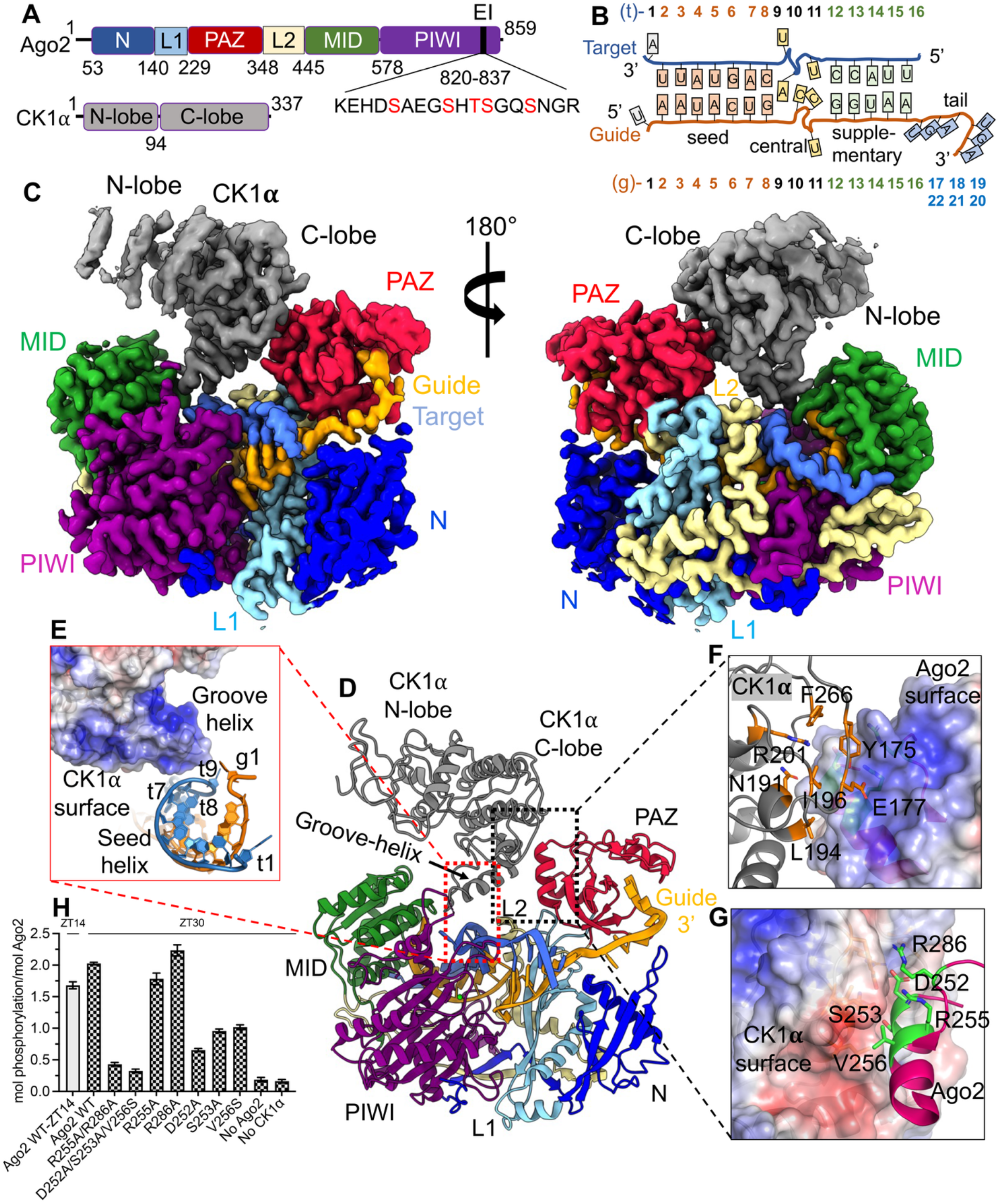
Cryo-EM structure of RISC^ZT30^ bound to CK1α kinase. (A) The domain architecture of Ago2 and CK1α, with the sequence of EI highlighting the conserved phosphorylation sites (red). (B) A schematic of the guide-target duplex used in the study. (C) Cryo-EM map for the RISC^ZT30^-CK1α complex in two different orientations and (D) a cartoon representation of the structure. (E) A zoomed-in view of the CK1α groove helix-RNA interface. The CK1α-Ago2 PAZ interface with (F) PAZ shown as surface and interacting CK1α residues shown as sticks and (G) vice-versa. (H) *In vitro* Ago2 phosphorylation with ZT30 targets with different PAZ mutations.

Ago2 displays the familiar canonical duck shape with the PAZ-domain sitting over the other three domains^10^. CK1α kinase docks onto the PAZ domain via its C-lobe, establishing a direct protein-protein interface burying ∼325 Å^2^ surface area. The kinase N-lobe is positioned above the Ago2 RNA-binding channel with the kinase catalytic site facing towards the MID-PIWI domain (**Fig 1D**). Density for AMPPNP was not observed, and only a partial segment of the EI could be built into the structure, which points towards the kinase catalytic site (**Fig 1C-D**). In this conformation, EI could reach the CK1α catalytic site, but is disordered, likely due to lack of interactions. Moreover, a 3D variability (3DV) analysis for RISC^ZT30^-CK1α shows slight motion of the PAZ and CK1α, moving as a unit towards the Mid domain across the RNA binding channel, whereas the EI peptide remains unresolved (**Supplementary movie 1-2**). A positively charged α6-helix (from the CK1α C-lobe) is positioned right above the seed-helix and interacts with the t7-t8 phosphates via its cQ231 and cK235 side chains (‘c’ refers to CK1α residues), sensing the presence of the target RNA (**Fig 1E, Supplementary Fig 2A**). At the protein-protein interface, the CK1α C-lobe and the Ago2 PAZ domain exhibit negative and positively charged complementary surfaces, and establish salt-bridge interactions (cE177-aR255, cR201-aD252 (‘a’ refers to Ago2 residues), polar interactions (cY175-aR255, cN191-aS253), and hydrophobic interactions (cL194-aV256) (**Fig 1F-G**). Mutating these Ago2 PAZ residues in clusters (R255A/R286A and D252A/S253A/V256S) almost completely ablates CK1α-mediated Ago2 phosphorylation with the ZT30 target RNA (**Fig 1H**), but shows a modest effect on guide or target binding (**Supplementary Fig 2B-C**). In addition, Ago2 D252A, S253A, or V256S point mutants but not the R255A or R286A mutants, reduced Ago2 phosphorylation with ZT30 (**Fig 1H**). Corresponding CK1α cluster mutants also reduced Ago2 phosphorylation with ZT30 (**Supplementary Fig 2D**). This mutational analysis underscores the importance of these PAZ and CK1α residues in establishing the protein-protein interface and promoting EI phosphorylation. In addition, during analytical size-exclusion chromatography (aSEC) analysis, a slightly smaller amount of mutant CK1α proteins co-eluted with RISC^ZT30^ compared to WT CK1α (**Supplementary Fig 2E**), consistent with weaker binding leading to reduced Ago2 phosphorylation. Interestingly, these residues are highly conserved in the PAZ domain of all four human Agos, but not in the PAZ domain of Piwi proteins (Piwi1, Piwi2, PiwiL4) (**Supplementary Fig 2F**), consistent with the absence of an EI and a phosphorylation-cycle in Piwi proteins. These PAZ residues are completely conserved in fly Ago1, which specializes in miRNA-mediated silencing, but poorly conserved in the slicing fly Ago2, which specializes in siRNA-mediated silencing (**Supplementary Fig 2F**) and also lacks the conserved phosphorylation sites in EI. This strongly suggests a conserved role for this interface, specifically in miRNA-mediated repression.

### A complete structural view of miRNA-target binding in Ago

Despite several structural studies, a complete miRNA-target duplex or even a complete miRNA guide has not been reported for Ago2. We now observed the full 22-nt miR200 guide bound in RISC^ZT30^-CK1α (**Fig 1D, 2A**) and RISC^ZT30^ structures (**Supplementary Fig 1F and Supplementary Fig 3A**), which exhibit a fully base-paired seed (g2-g8), a structured untwisted central (g9-g11), a fully base-paired supplementary (g12-g16), and a structured tail (g17-g22) region. The target t2-t8, t12-t16 nucleotides base pair to form the seed-helix and supplementary-helix respectively, t9-t11 reside in the central channel (**Fig 2B and Supplementary Fig 3B**), while t1-A fits into a conserved pocket^12,26^ at the MID-PIWI interface. The 5’ region of the target (t17-t30) is not observed in the structure and remains solvent exposed. The RNA exhibits several stabilizing interactions with Ago2, which translate into distinct conformational changes in particular protein elements, as described below.

**Figure-2.**
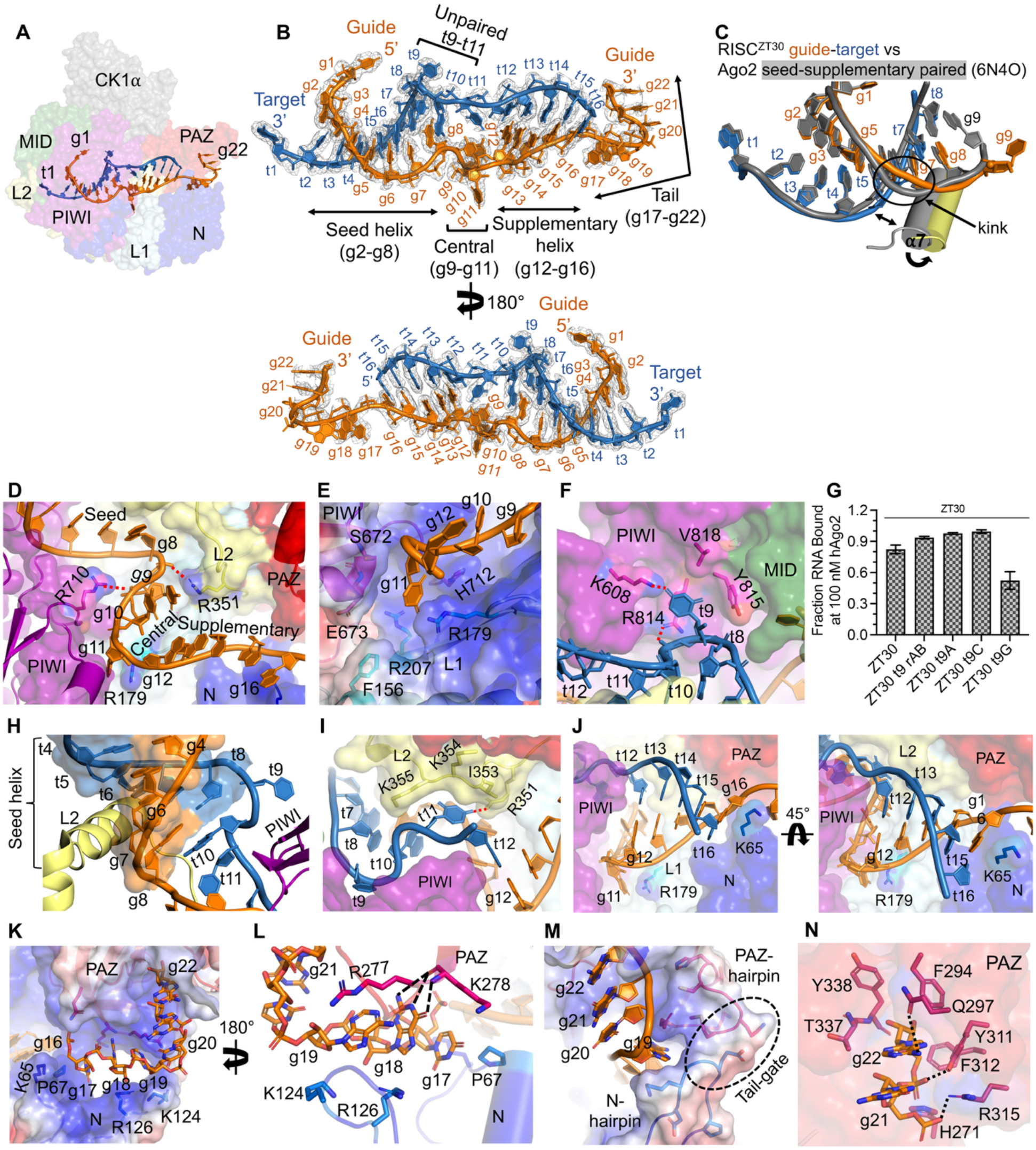
The miRNA guide-target duplex in the RISC-silencing complex. (A) The cryo-EM map with the complete guide-target complex observed in the RISC^ZT30^ structure highlighted. (B) Observed cryo-EM map density of the RNA shown in two orientations. The seed-helix, unpaired central duplex, supplementary-helix, and single-stranded tail regions are marked. (C) A zoomed-in view of the guide g6-g7 kink. The kink in the seed-supplementary paired structure (gray) straightens out in the RISC^ZT30^-CK1α structure (gold) with a corresponding repositioning of the L2 α7 helix. (D) The distorted geometry of the central region passing through the narrow central channel in Ago2. R351:g9-g10 are stacked. (E) g11 flips into the g11 pocket between PIWI-L1 and stacks on R179. The backbone has a significant bend at g11-g12. (F) The t9 pocket in PIWI showing interactions with t9. (G) *In vitro* Ago2 phosphorylation with t9 variants of ZT30 target showing reduced phosphorylation with t9-G substitution. (H) The t10 nucleotide stacks under the seed-helix (shown as surface) (I) t11 interactions with the L2 loops in the central channel. t12 is positioned in the supplementary chamber base-pairing with g12. (J) The full supplementary helix shown in two orientations. R179-g11 and K65-g16 stacking support positioning of the supplementary helix. (K) The tail region (sticks) passes through the positively charged N-PAZ channel and the PAZ 3’ binding pocket. (L) The interactions of g17-g19 in the N-PAZ channel. (M) The tail bending into the PAZ domain assisted by the structured N-PAZ hairpins; the tail-gate (surface) and (N) interactions of g21-g22 in the PAZ 3’-binding pocket. Additional interactions observed are shown with black dotted lines.

### The Seed helix

The g2-g5 sub-seed region, used in initial target screening, is structurally conserved. However, the g6-g8 seed region is smoother and no longer exhibits a backbone kink as observed in previous structures^19,20,27^ (**Fig 2C**). The L2 loop in those structures is positioned slightly towards the seed; more specifically near the g8 nucleotide. We observe, however, that in the RISC^ZT30^-CK1α structure, L2 opens just enough so g8 slides under the L2 loop, releasing the tension at g6 and smoothing the g6-g8 backbone kink. Moreover, smoothing out the g6-g8 seed slightly alters the local geometry around g5-8/t5-8, translating into a slightly more open L2 helix-7 (**Fig 2C**). Following g8, the RNA backbone turns away from L2 towards the PIWI domain, placing the g9 nucleotide in the central chamber. Nucleotides t2-t8 base pair with the seed forming an A-form double-stranded seed-helix, stabilized by the L2 helix-7 (**Supplementary Fig 2A**).

### The central RNA region

The conformation of the RNA in the central region is the most surprising feature of this structure, since it is pried apart by the protein, interrupting the helical geometry of the seed-helix, so that the two strands are no longer intertwined in the central region. The three unpaired guide and three target nucleotides are constrained within the narrow central chamber. Right after the seed, the guide backbone turns almost 180°, directing the g9-g11 towards the PIWI domain. The base of g9 stacks with R351 (L2) side chain on one side and g10 on the other, and its sugar lands in a positive pocket in the L1 β-sheet, supporting the major distortion in the backbone between g8 and g9 (**Fig 2D**). The g9 base in previous Ago2 structures that contain short target RNAs is stacked with the last seed-pair (ie:g8/t8)^19,20^, but with a complete structured miRNA-target, Ago2 pulls the g9 sugar into a previously unrecognized pocket (**Fig 2D**).

As mentioned, the g9 and g10 nucleotide bases are stably stacked on each other, and their WC edges point towards the supplementary region. However, g11 flips and points into a deep positively charged pocket formed between PIWI and L1 domains (**Fig 2D-E**). This previously unrecognized pocket is carved between R179 (from L1) and the S672-E673 loop (PIWI), which anchors the g11 nucleotide and supports a sharp turn in the RNA backbone between g11 and g12, directing the RNA that follows towards the N domain (**Fig 2E**). Ago2 R179A exhibits slow miR200 guide binding and a ∼3-fold reduced binding affinity (**Supplementary Fig 3C**), consistent with a role in stabilizing the unwound guide. Moreover, we tested two additional miRNA-target pairs (miR132-TJAP1 and miR1-SOX9). Together with miR200-ZT30, all three cases exhibited a similar target-binding-triggered Ago2 phosphorylation **(Supplementary Fig 3D)**, indicating the same EI-phosphorylation-mediated target release mechanism. The R179A mutation resulted in slower target binding **(Supplementary Fig 3E-G)** and a modest reduction in binding affinity **(Supplementary Fig 3H)** for the 30-nt long ZT30, TJAP1 and SOX9 targets, relative to their corresponding WT-RISCs. The observation strongly suggests that R179-mediated support of an untwisted central region is likely a general feature in miRNA-RISC for these types of guide-target pairs. The structural comparison between guide-bound^15^, guide-seed paired^19^, guide-seed+supplementary paired^20^, and our RISC^ZT30^ structure suggests that the g11 pockets are a pre-existing feature in Ago2, but require a slight opening of N and L1 domains away from PAZ (widening of N-PAZ channel) to accommodate the g11 nucleotide adequately (**Supplementary Fig 3I**).

On the target strand, the t9 nucleotide is flipped and placed into a positively charged pocket in the PIWI domain. R814 stacks with t9-U and interacts with t9 phosphate via an H-bond, whereas K608 H-bonds with t9-U(O4). Additionally, Y815 and V818 line the pocket without any direct interactions with t9 (**Fig 2F**). Although U seems to be specifically recognized in the t9 pocket by K608, a guanine base can fit with minor rearrangements (in R814), as shown previously^28^ (**Supplementary Fig 3J**). Interestingly, a t9-G substitution, but not t9-A, t9-C, or t9 abasic substitutions, reduces ZT30 target binding (fraction RNA bound at Ago2 saturation) in RISC (**Fig 2G and Supplementary Fig 3K**). This feature might be helpful for fast target release from RISC after recruiting the downstream factors in humans, as also suggested for HCV target RNAs, which have a conserved t9-G throughout^28^. However, the phosphorylation level does not change for t9-substituted ZT30 RNA-loaded RISC (**Supplementary Fig 3L**).

This region of the target (t9-t11) is clearly observed in these structures. Following the unexpected flip of t9, the RNA backbone at t10 turns almost 180° from PIWI towards the L2 loop and the base of t10 stacks under the last base-pair of the seed-helix (g8-t8), stabilizing the seed-helix end (**Fig 2H**). However, RISC target binding with an abasic t10 is only modestly affected (**Supplementary Fig 4A-B**). In contrast, Ago2 structures containing an unstructured target central region^19,20,28^ show that g9 could stack under the seed-helix as well (**Supplementary Fig 4C**). Taken together, it seems that anchoring the seed-helix end is a conserved feature in RISC, which can be executed by g9 or t10 when g9 is pushed into an L2 pocket (see above).

At the t11 nucleotide, once again the RNA backbone turns prominently from L2 towards the supplementary chamber, and tucks t11-U under the L2 loop (at I353-K354-K355) with U(O4) H-bonding with L2 loop R351 (**Fig 2I**), which, in turn, also stacks on g9; see **Fig 2D**). The L2 loop (351-358) and PIWI loop (602-607) are known key features of the narrowed central chamber, preventing central pairing^20^. This chamber opens up to facilitate the target rolling around the guide to assemble the slicing competent state^29^. Our structure highlights that these loops also support the guide-target central region, albeit with their slight repositioning (**Supplementary Fig 4D**). Overall, the constricted central region is highly distorted keeping the two strands from intertwining, which could facilitate target removal when the target is not sliced.

### The supplementary helix

The A-form supplementary helix sits in the Ago2 supplementary chamber. Nucleotides g12-g16 interact with L1 and N-domain via their backbone, while t12-t16 do not interact with the protein (**Fig 2J**). A previous study suggested that a clash with the L2 loop would not allow t12 to base pair with the guide^20^, but our structure shows that with slight repositioning of the L2 loop (see **Supplementary Fig 4D**), Ago2 accommodates a fully five base-paired supplementary helix. Notably, Ago2 anchors the guide supplementary region (g12-g16) by stacking R179 (L1) and K65 (N-domain) against g11 and g16, respectively (**Fig 2J**). An R179A or K65A mutation in Ago2 reduces guide binding by ∼3 fold and ∼5 fold, respectively (**Supplementary Fig 3C**), confirming their role in stabilizing the supplementary helix. Both R179 and K65 are conserved in all Agos and seem capable of stacking with any nucleobase at these positions, suggesting that it could be a common mechanism to support the guide supplementary region.

### The tail

The guide tail (g17-g22) stays unpaired with the 3’ end nestled in the PAZ domain, while the target is not observed after t16 due to a lack of interactions with RISC or the guide. The stacked g17-g19 nucleotides establish extensive interactions with Ago2 as they pass through a positively charged, wide open N-PAZ channel with their bases pointing into the channel (**Fig 2K**). The g17 nucleobase is pushed against P67 from a repositioned α1-helix in the N-domain (**Fig 2L**), whereas the g18-G nucleobase stacks on the guanidinium group of R126 (N-domain) and H-bonds with K278 main chain carbonyl and amide (PAZ). The g19 base stacks between conserved R277 (PAZ) and K124 (N-domain), supporting the bend in the tail that follows (**Fig 2L**). In this conformation, any base substitution at g17 and g19 position, would easily fit and establish similar interactions, suggesting a common mode of tail binding in RISC.

The bend in the backbone at g19-g20 is supported by structured hairpins from the N-domain (aa 120-125) and PAZ domain (aa 270-276) at the end of the N-PAZ channel, which we call the ‘tail-gate’. This gate directs the stacked g20-g22 tail into the 3’ binding pocket in the PAZ domain (**Fig 2M**). Nucleotides g21-g22 are completely buried in the PAZ domain and in addition to the previously observed conserved PAZ interactions (**Supplementary Fig 4E**) g22 base interacts with Q297, g22 phosphate withY311, and g21 phosphate with R315 (**Fig 2N**).

### Supplementary pairing induces CK1α favorable RISC rearrangements

Our previous study showed that CK1α-mediated Ago2 phosphorylation drastically increases as base pairing in the supplementary region progresses from 13 to 14 nucleotides during target binding^6^. Therefore, we investigated the role of supplementary pairing by determining single-particle cryo-EM structures of RISC^ZT13^ and RISC^ZT14^ containing a 13-nt and 14-nt ZT1 target RNA, respectively, in the presence of CK1α (**Fig 3**). For RISC^ZT13^ we observed particles in two predominant CK1α-free states (denoted as RISC^ZT13^) (**Supplementary Fig 5A**), and generated RISC^ZT13^-closed (∼3.4 Å) and RISC^ZT13^-open (∼3.7 Å) cryo-EM maps (**Supplementary Fig 5B-C**). A subset of particles from RISC^ZT13^ also produced a cryo-EM map with much weaker density corresponding to traces of CK1α (RISC^ZT13^-CK1α-bound), but could not be confidently built (**Supplementary Fig 5A, S5D**). However, for RISC^ZT14^ we observed about 65:35% particle distribution in RISC^ZT14^ and RISC^ZT14^-CK1α states, producing ∼3.4 Å and ∼3.7 Å resolution cryo-EM maps, respectively (**Supplementary Fig 5E-G**). We used the RISC^ZT30^ structure for model building. Final statistics are summarized in **Supplementary Table 1**.

**Figure-3.**
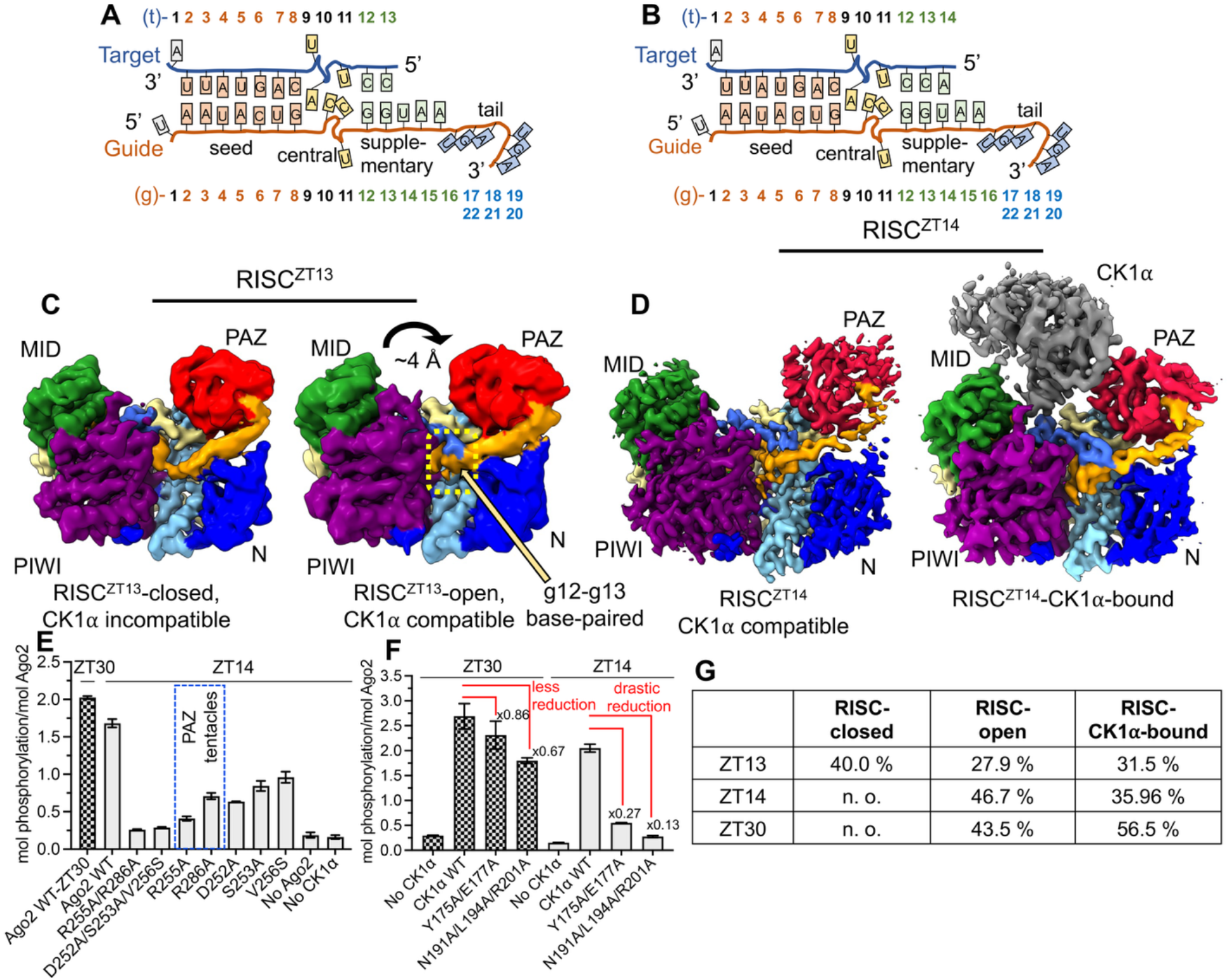
RISC dynamics upon supplementary base-pairing. A cartoon representation of the guide-target duplex used for (A) RISC^ZT13^ and (B) RISC^ZT14^ analysis. (C) Cryo-EM maps for RISC^ZT13^ in closed (left) and open (right) conformations. The movements in the PAZ and base-paired g12-g13 region are indicated. (D) Cryo-EM maps for RISC^ZT14^ in the CK1α-free (left) and CK1α-bound (right) states. t14 is clearly base-paired in both maps and shows CK1α-compatible PAZ positioning in both maps. (E) *In vitro* Ago2 phosphorylation of different PAZ mutations with ZT14 and (F) CK1α cluster mutations with ZT30 (checkered bar) and ZT14 (grey bar). Reduced phosphorylation levels in different conditions are marked. (G) The calculated no. of cryo-EM particles used in different 3D map generations for the structures presented in this study.

RISC^ZT13^-closed contains ZT13 target density up to t10 with only seed pairing visible, while RISC^ZT13^-open state clearly exhibits density for the complete ZT13 target base-paired up to nucleotide g13 in the supplementary region (**Fig 3C**). The RISC^ZT13^-open structure is almost identical to the RISC^ZT30^ structure with an RMSD of 0.4 Å (**Supplementary Fig 6A-B**). Interestingly, with the g12-g13 base pairing, the RISC^ZT13^-open structure exhibits a ∼4 Å movement of the PAZ away from the MID domain, bringing the bound g20-g22 tail with it (**Fig 3C**), which widens the exposed RNA-binding channel. Notably, the structural superposition also shows that CK1α would clash with the PAZ domain in RISC^ZT13^-closed, whereas the open conformation is better suited for CK1α docking (**Supplementary Fig 6C-D**).

The RISC^ZT14^ structure in both RISC^ZT14^ and RISC^ZT14^-CK1α (**Fig 3B,3D**) is almost identical, and the two structures superimpose well with RISC^ZT30^ (**Supplementary Fig 6E-F**). In RISC^ZT14^, the CK1α groove helix interactions with the target are conserved with RISC^ZT30^. The CK1α Q231A/K235A mutant reduces Ago2 phosphorylation with ZT14 and ZT13 targets but not with the ZT30 target (**Supplementary Fig 6G**), consistent with their role in sensing the seed helix geometry and EI phosphorylation during initial base pairing of the supplementary region. The CK1α-Ago2 PAZ interface interactions (**Supplementary Fig 6H**) are conserved with RISC^ZT30^-CK1α as well. A 3DV analysis of RISC^ZT14^-CK1α (**Supplementary movie 3-4**) shows a similar motion of CK1α and PAZ towards the Mid domain as seen for RISC^ZT30^-CK1α (**Supplementary movie 3-4**) Cluster mutations of PAZ domain residues at this interface (aR255A/R286A and aD252A/S253A/V256S) or CK1α (cY175A/E177A and cN191A/L194A/R201A) drastically reduced Ago2 phosphorylation in RISC^ZT14^ (**Fig 3E, 3F**). Interestingly, the effect of CK1α mutations on Ago2 phosphorylation is significantly more pronounced (up to 4-fold higher) with a 14-nt target (RISC^ZT14^) than with a 30-nt target in RISC^ZT30^ (**Fig 3F**), suggesting a greater contribution of this interface when only half of the supplementary region is base paired as in RISC^ZT14^. Furthermore, unlike in RISC^ZT30^ (see **Fig 1G**), all the PAZ point mutations at that interface, including aR255A and aR286A, drastically reduced Ago2 phosphorylation in RISC^ZT14^, suggesting that this interface is more sensitive to mutations when only half of the supplementary region is base paired. Considering these biochemical observations, it is possible that the state exemplified by RISC^ZT14^ acts a checkpoint in CK1α loading, which once progressed with further base pairing would lead to hierarchical EI phosphorylation, thereby showing a greater effect on phosphorylation. Similarly, all the CK1α point mutations at this interface reduced Ago2 phosphorylation in RISC^ZT14^ (**Supplementary Fig 6I**) and not in RISC^ZT30^ (see **Supplementary Fig 2D**), underscoring the dynamic role these residues have at a specific step in kinase binding to Ago2.

The guide-target geometry in RISC^ZT14^ is conserved with RISC^ZT30^, but in the absence of CK1α docking onto RISC^ZT14^, the PAZ domain appears to be more dynamic based on a 3DV analysis. Here, PAZ movements towards the Mid domain of RISC^ZT14^ are correlated with a disappearance of cryo-EM density for the central region (**Supplementary movie 5**). In keeping with this, PAZ movement is similar to that in the RISC^ZT13^-closed state, which also lacks a structured central region. Moreover, unlike with a ZT30 target, substitutions at central t9 do not affect ZT14 target binding in RISC (**Supplementary Fig 6J-K**) or Ago2 phosphorylation by CK1α (**Supplementary Fig 3L**). Interestingly, an abasic t10 nucleotide containing ZT14 (ZT14-t10-rAb) but not ZT30 (ZT30-t10-rAb) target RNA exhibits significantly (∼38%) reduced Ago2 phosphorylation (**Supplementary Fig 6L**), suggesting that g8/t8 stacking by the kinked t10 is important when only half of the supplementary region is paired, though RISC target binding with an abasic t10 is only modestly affected (**Supplementary Fig 4A-B**). These effects appear to be compensated by complete supplementary region pairing, which renders the complex more robust to protein mutations and RNA alterations.

Notably, the proportion of cryo-EM particles contributing to the different structures indicates that the RISC^ZT13^-closed form is predominant over the RISC^ZT13^-open state, with about 31% particles converging into a weak RISC^ZT13^-CK1α complex. However, with one more nucleotide in the target as is the case with ZT14 binding, not only does the closed state completely disappear, but the particle distribution of the CK1α-bound state and the CK1α map quality significantly improve, strongly suggesting that the additional pairing of g14 supports CK1α binding to Argonaute. With further pairing in the supplementary region (RISC^ZT30^), this number increases to ∼ 56% particles in a CK1α-bound state (**Fig 3G**). Taken together, supplementary pairing in g14 and PAZ repositioning are interlinked and permit CK1α binding, resulting in increased Ago2 phosphorylation in RISC^ZT14^ compared to RISC^ZT13^.

### CK1α anionic binding site supports Ago2 EI phosphorylation

Although the EI was not visible in the cryo-EM maps, we investigated how CK1α could bind the EI peptide during hierarchical phosphorylation. Analysis of the CK1δ structure in complex with tungstate (PDB 1CKJ)^30^ or with phosphorylated p63 peptides (PDB 6RU8)^31^ pointed toward two positively charged conserved anionic binding sites (ABS) (**Fig 4A**) in CK1α formed by R186-G223-K232 and R135 (**Fig 4B-C**), which might accommodate a phospho-serine in EI. Moreover, an AlphaFold 3.0^32^ structure prediction of RISC^ZT30^ containing phosphorylated Ago2 on S824, S282 and S831 shows the phospho-S828 coordinated in the CK1α catalytic site, while phospho-S824 and S831 are stabilized by interactions within the CK1α ABS pockets (**Fig 4D**).

**Figure-4.**
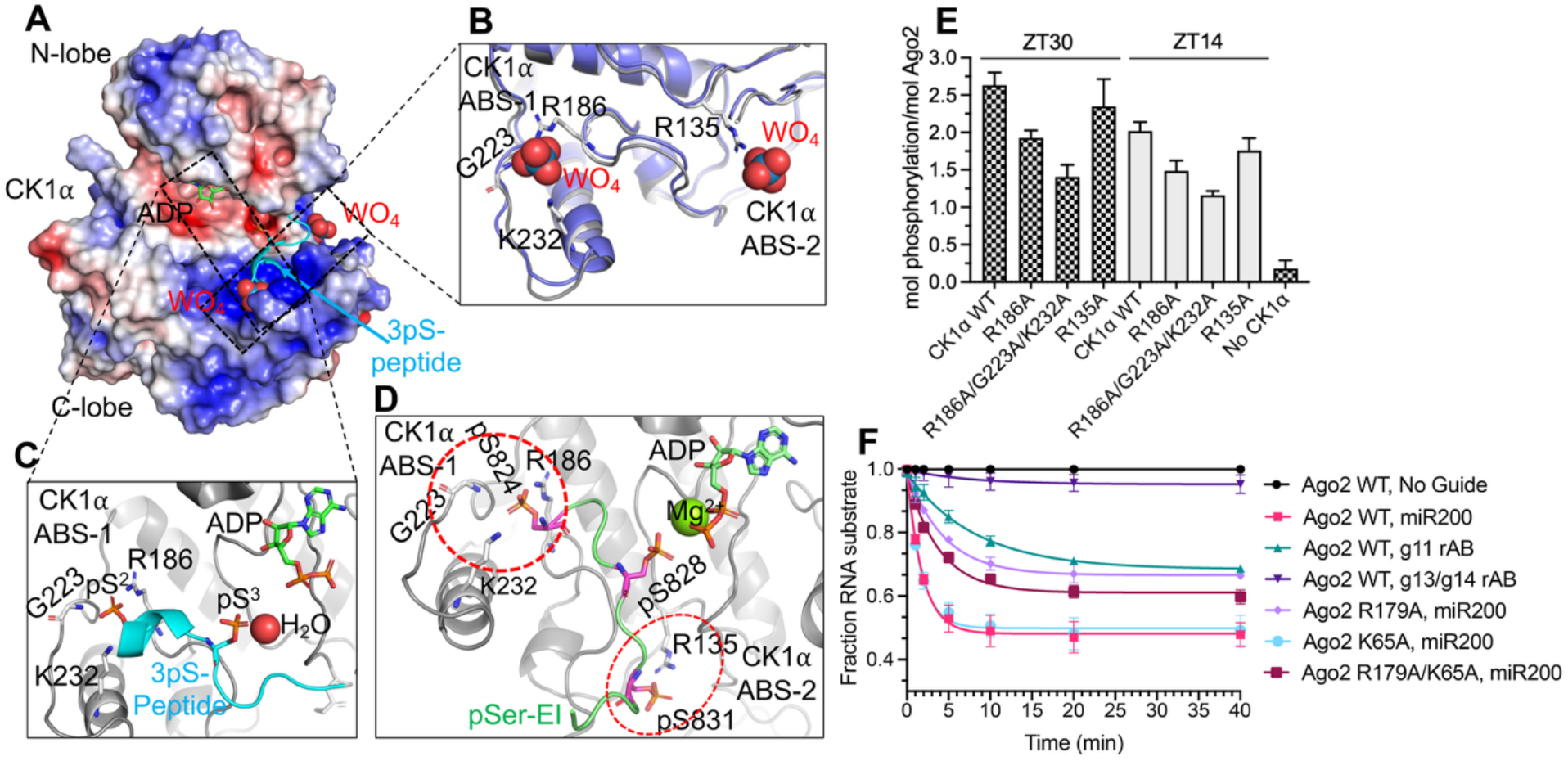
The CK1α ABS supports Ago2 EI phosphorylation. (A) The electrostatic surface of the CK1α from RISC^ZT30^, showing the positively charged ABS occupied with superimposed tungstate ions from CK1δ (PDB 1CKJ) and CK1δ-3pSer-peptide complex (cyan) (PDB 6RU8) crystal structures. (B) A zoomed-in view of the tungstate binding site. (C) The 3pSer-peptide binding in the ABS sites. (D) An AlphaFold 3.0 model prediction of the CK1α ABS sites bound with Ago2 EI pS824 and pS831. pS828 is seen bound in a post-catalytic state in the active site. (E) *In vitro* phosphorylation of Ago2 with different CK1α ABS mutants with ZT30 (checkered bar) and ZT14 (grey bar) targets. (F) The target RNA slicing assay using different Ago2 mutants and guide variants as indicated.

To validate the role of CK1α ABS, we tested CK1α R186A, R186A/G223A/K232A, and R135A mutants in phosphorylation assays of Ago2 with either ZT30 or ZT14 target RNAs. The CK1α R186A mutation clearly reduced Ago2 phosphorylation, while the R186/G223A/K232A mutation had an even larger effect with both ZT30 and ZT14 target RNAs (**Fig 4E**), confirming the role of CK1α ABS in efficient Ago2 phosphorylation. However, the CK1α R135A mutation only modestly reduced Ago2 phosphorylation with ZT14, but not with ZT30 as the target RNA, suggesting this site’s involvement during initial supplementary pairing steps. Notably, these ABS mutants do not abolish CK1α binding to RISC^ZT30^, and co-elute with RISC^ZT30^, similar to the WT CK1α, as analyzed by aSEC (**Supplementary Fig 7A**), suggesting that the observed reduction in phosphorylation is due to impaired EI binding in ABS but not overall binding of CK1α to RISC^ZT30^.

### RISC slicing-competent state requires central g11 stacking

RISC adopts a slicing-competent state upon encountering a target RNA fully complementary to the guide. This catalytic state was suggested to arise through a two-helix state (RISC with a seed-helix and supplementary helix)^33^, and involves rolling of the supplementary helix over the N domain to form a double helix in the central chamber, which releases the 3’ guide from the PAZ^29,34^. Since our RISC^ZT30^ structure could represent a two-helix state of sorts, we analyzed the role of the new features we observed in the RISC^ZT30^ structure in slicing of a fully complementary target. Notably, mutation in the g11-stacking R179 residue (see **Fig 2E**) leads to significantly impaired slicing of a fully complementary target as compared to WT-Ago2. Concurrently, using a miR200b guide with an abasic nucleotide at g11 also impairs target slicing with reduced product accumulation (**Fig 4F**). The pocket formed by R179 (L1) in the two-helix state disappears as the N-L1 domain opens and RISC transitions to the slicing state. Therefore, R179-g11 interactions likely occur transiently and precede the RISC slicing state. Mutation of K65 that stacks with g16 (see **Fig 2J**) shows no effect on target slicing, and R179A/K65A double mutant exhibits reduced slicing levels similar to the R179A point mutant (**Fig 4F**), suggesting that K65 or g16 stacking is not a critical slicing determinant. Furthermore, disturbing the supplementary helix formation using a miR200b guide with abasic g13-g14 nucleotides abrogates target slicing, confirming that a two-helix state is a prerequisite for target slicing by Ago2.

### Structural Dynamics during CK1α docking

A comparative analysis of Ago2 structures revealed major structural dynamics affecting the N-PAZ channel, guide, and supplementary-helix binding. In the Ago2 structure containing a paired seed+supplementary region (PDB 6N4O)^20^, the N-PAZ channel is narrow, with the tail-gate and other PAZ loops (aa 296-301 and 332-334) unstructured. In the RISC^ZT30^ structure, the PAZ domain rotates/rolls towards the guide and moves ∼5 Å towards the N-domain hairpin (**Fig 5A-B**). Concurrently, the N and L1 domains open up 5 Å and 2.5 Å, respectively, pointing away from the MID domain (**Fig 5C**), effectively widening the N-PAZ channel. The repositioned PAZ domain also pushes the bound tail by 6-10 Å towards the N-domain and the whole supplementary helix in Ago2 (**Fig 5D**). The tail-gate hairpins move to completely close the wider N-PAZ channel supporting the guide tail (**see Fig 2M**). These acrobatics allow Ago2 N-PAZ to comfortably fit 3 tail nucleotides (g17-19) (**Fig 5E**), while only 2 nucleotides are observed in the Ago2 structure with a paired seed+supplementary region (**Fig 5F**) that has a wide open tail-gate (**Supplementary Fig 4F**). This widening of the N-PAZ channel might be a mechanism by which Ago2 can accommodate different miRNA guide lengths.

**Figure-5.**
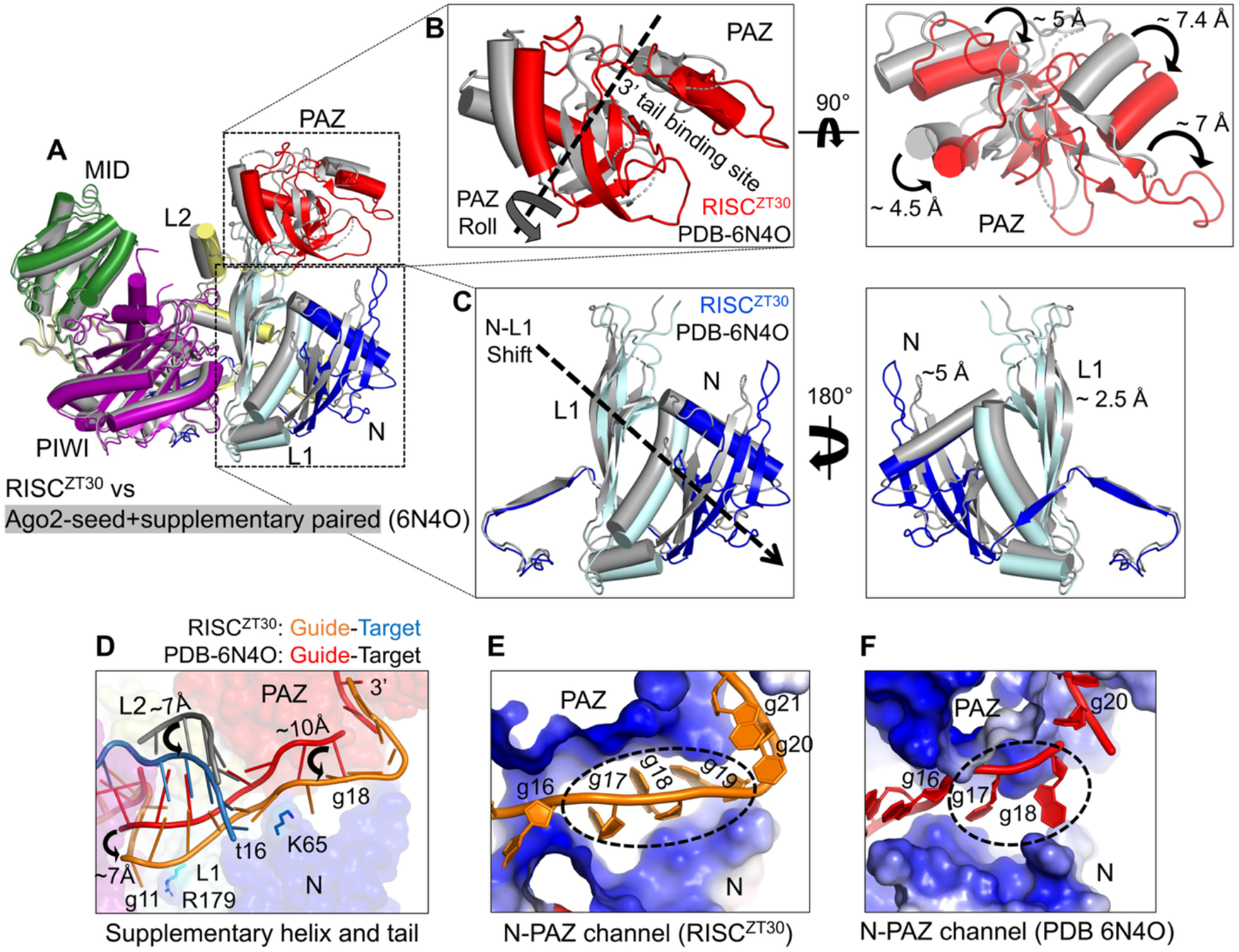
Structural dynamics in RISC. (A) Ago2 superposition of RISC^ZT30^-CK1α (colored) and the Ago2 crystal structure with paired seed+supplementary region (in grey) (PDB 6N4O). (B) A close-up view of the PAZ domain in two orientations. (C) N-L1 domains shown in two orientations. The displacements in the structural elements are marked. (D) The difference in the positioning of the supplementary-helix and tail region in RISC structures. The displacements of the different RNA segments are marked. (E) The N-PAZ channel as observed in RISC^ZT30^ (right) and Ago2 crystal structure (left) showing bound tail nucleotides.

## Discussion

Ago2 phosphorylation-mediated target release is a conserved and widely used mechanism for efficient RNAi across different species^4,5^, but the molecular mechanism underlying this process remained unknown. Our study reveals a stepwise mechanism for CK1α docking onto RISC as the guide RNA’s supplementary region progressively base pairs with the target RNA (**Fig 6**). Ago2 PAZ is the most dynamic domain in the RISC core and holds onto the 3’ end tail. Upon RISC’s encounter with a target, a double-stranded seed helix is assembled, while the supplementary region (g12-g16) stays unpaired. Ago2 stays in a closed conformation, with the PAZ domain closer to the MID domain, unfavorable for CK1α binding, resulting in a narrow central RNA-binding channel in the absence of supplementary base pairing^15,19^. The flexibility of the PAZ domain appears to be influenced by supplementary base pairing as the PAZ swings away from the MID domain to an open, CK1α-compatible conformation as g12-g13 nucleates the supplementary helix. Unlike the slicing state, the PAZ movement in the silencing states is minor and does not involve release of the 3’ tail. The more open PAZ at this stage (RISC^ZT13^) is compatible with CK1α docking and exhibits only minimal Ago2 phosphorylation, likely due to the inherent flexibility of the PAZ. Further base pairing of g14, resulting in a three base-paired supplementary RISC-target state (RISC^ZT14^), restricts the PAZ exclusively to the open conformation, which facilitates CK1α docking and establishes a protein-protein interface with the PAZ via its C-lobe, and a protein-seed helix interface via its α6-helix. Thus, improved access to CK1α effectively explains the jump in Ago2 phosphorylation levels with a 14-mer target as in RISC^ZT14^. This observation also supports the idea that 14 nt represents the shortest target length capable of forming the initial stable supplementary pairing. In the next steps, the full supplementary region base pairs and RISC attains a two-helix state (RISC^ZT30^). Though CK1α interactions with PAZ do not change anymore, increased Ago2 phosphorylation at this stage is attributed to longer retention times of CK1α on RISC, resulting in hierarchical phosphorylation of the EI, supported by the ABS sites in CK1α, ultimately leading to target RNA release. The conserved Ago EI phosphorylation sites correlate with a conserved CK1α-PAZ interface among different species, effectively pointing to conserved structural dynamics and a molecular mechanism for optimized miRNA-mediated repression across species.

**Figure-6.**
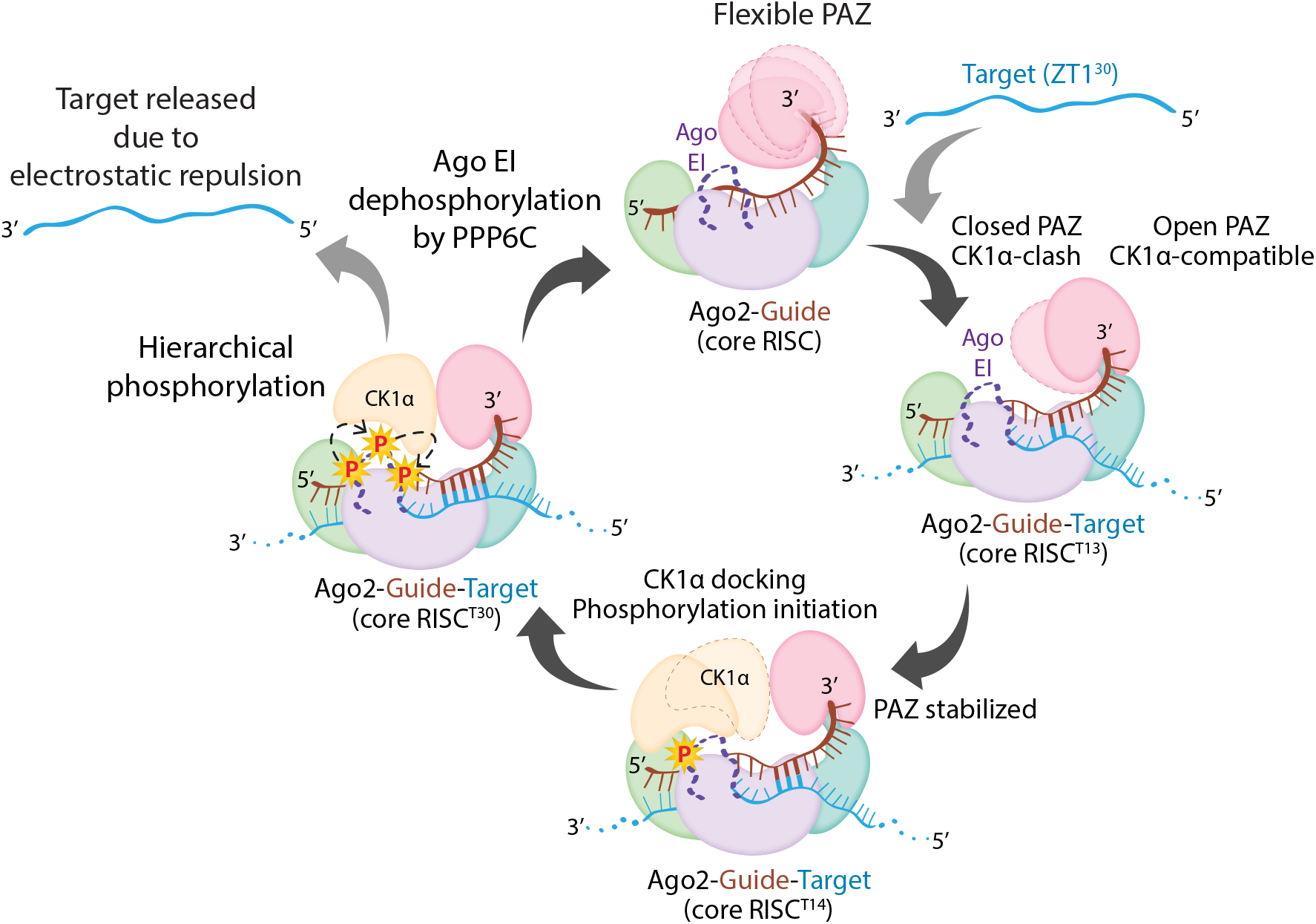
Structural model for supplementary pairing-triggered Ago2 phosphorylation by CK1α. PAZ in the RISC core is flexible prior to target binding. Initial base-pairing with the target in the supplementary region starts to restrict the PAZ domain positioning (as in RISC^ZT13^). Once g14 is based paired with its target (RISC^ZT14^), the PAZ domain is restricted to an open conformation, facilitating initial CK1α binding and subsequent EI phosphorylation. Full supplementary region pairing with the target (as in RISC^ZT30^) stabilizes the complex and further increases EI phosphorylation, leading to target release and RISC recycling. The target does not twist or intertwine over the guide in the central region, and the duplex is held that way by Ago2. This enables the target to detach from the complex more easily when slicing does not occur.

Ofcourse, a possible influence of additional silencing factors, such as the deadenylase and decapping complexes, brought in by TNRC6A (or GW182 in flies) cannot be ruled out without further investigation. Among the miRNA binding sites in human 3’ UTRs, a few thousand, accounting for about 5% of target sites, require base-pairing of the 3’ supplementary region for silencing^1,35^. Formation of the supplementary helix increases the RISC-target affinity by up to 500-fold^36^, which necessitates an active mechanism for effective target release. Our study shows that the formation of the 3’ supplementary helix allows for CK1α-mediated phosphorylation of Ago2 for several guide-target pairs *in-vitro* (to enable RISC recycling (employing charge repulsion to overcome a higher guide:target affinity), Since RNA helix assembly does not strictly require perfect Watson-Crick base-pairing^37–39^, a 3’ supplementary helix could form with a number of mismatched base-pairs, suggesting that a similar mechanism for Ago2 EI phosphorylation could be at play at a majority of these types of target sites.

Our study also produced a complete structural view of a miRNA-target duplex, providing new insights into several key aspects of Ago2 miRNA-mediated silencing. Most notable is that the guide and target do not intertwine in the central region, with the protein pulling apart bases to untwist the RNA duplex. This would greatly enable the target’s release post silencing in the absence of a slicing event at the center of this region. It was always perplexing how the target would detach from the complex. But with a target that is much less intertwined with the guide, coupled with phosphorylation of the EI leading to charge repulsion with the target, we uncovered a mechanism for effective target release. Other features include: straightening of the g6-g7 kink; t10 stacking on the seed-helix and t11-L2 loop repositioning; book-ending the supplementary helix by R179 and K65 on both ends; widening of the N-PAZ channel; and, finally, a structured tail-gate that directs the 3’ end of the guide into the PAZ domain.

Provided that target levels exceed cellular miRNA levels^40–42^, efficiency of target turnover determines the global miRNA-mediated repression. For the RISC silencing complex, this is achieved by target engagement, triggering Ago2 EI phosphorylation, electrostatically repelling the target^6^. With the central region untwisted, the repulsion would enable melting of both seed and supplementary helices without much unwinding, effectively increasing the target k_off_ by several fold, thereby releasing the target. Whereas in the RISC slicing complex, target cleavage between t10-t11 would release the helical tension, allowing unwinding of the RNA duplex. A widely opened N-L1 and PAZ domain channel loses most RNA interactions with the supplementary and tail helices respectively before cleavage^34^, supporting a rapid unwinding and target release.

Our observations that the supplementary helix and R179-g11 stacking affect slicing support the model where the slicing complex attains a two-helix or two-helix-like state^20,29,33^. In the slicing two-helix state, the position of the PAZ domain prevents CK1α docking, explaining why Ago2 phosphorylation is not triggered with a fully complementary target^6^. Furthermore, the twisted and untwisted conformations of the central guide-target duplex correlate with slicing for the former and silencing for the latter, which may also influence PAZ movements and thus CK1α compatibility.

## Materials & Methods

### Protein Expression and Purification

Recombinant HsAgo2 was prepared as previously described^15^, and dephosphorylated as described in Bibel et al 2022^6^, with minor modifications. Briefly, HsAgo2 was expressed with an N-terminal TEV protease cleavable Twin-Strep-SUMO fusion tag in Sf9 insect cells using the MultiBac system^43^. The insect cells were infected with baculovirus at 27°C for 60 hrs, harvested in Ago2 resuspension buffer (50 mM Tris pH 8.0, 100 mM KCl, 5 mM DTT supplemented with a protease inhibitor (PI) mix (pepstatin, leupeptin, PMSF, benzamidine, and aprotinin). KCl concentration was increased to 250 mM in resuspended cells before sonication. After Strep-Tactin affinity chromatography (Strep-Tactin 4Flow High Capacity, IBA) (50 mM Tris pH 8.0, 250 mM KCl, 5 mM DTT), the Twin-Strep-SUMO tag was cleaved by TEV protease overnight at 4°C. The RNA-free Ago2 was separated from Ago2 loaded with endogenous (insect) RNAs over a MonoS 10/300 or HiTrap SP-HP column (Cytiva) with a stepwise KCl gradient. RNA-free Ago2 was dephosphorylated by incubating overnight at 4°C with recombinantly expressed and purified λ protein phosphatase (λPP) at a 1:2.5 molar ratio, in the presence of 1 mM MnCl_2_. The dephosphorylated Ago2 was further purified via size-exclusion chromatography (SEC) on a Superdex 200 Increase 10/300 column (Cytiva) pre-equilibrated in 10 mM Tris pH 8.0, 200 mM KCl, 5 mM DTT, and 10% glycerol. Pure HsAgo2 containing fractions were concentrated and stored at −80°C. Different mutants of HsAgo2 were expressed and purified similarly to WT-HsAgo2. The HsAgo2 protein used for cryo-EM was purified similarly, apart from 25 mM HEPES pH 7.5 replacing Tris in the SEC buffer.

Human CK1α (isoform 1) was expressed in Sf9 or HighFive insect cells with an N-terminal TEV-cleavable Twin-Strep-SUMO tag using the MultiBac system. Following Strep-Tactin affinity chromatography (50 mM Tris pH 8.0, 200 mM NaCl, 5 mM DTT), the tag was removed by overnight TEV protease treatment and CK1α was further purified by cation exchange (HiTrap SP-HP, linear NaCl gradient elution) and size exclusion chromatography (Superdex 75 Increase 10/300 GL, Cytiva) in 10 mM Tris pH 8.0, 200 mM NaCl, 5 mM DTT. Pure protein fractions were concentrated, 20% glycerol added, and stored at –80°C. Different mutants of CK1α were expressed and purified similar to WT-CK1α.

λPP was cloned into a pET28 vector with an N-terminal 6xHis tag and expressed overnight at 18°C using IPTG in E. coli BL21(DE3) grown in TB media. λPP was purified by Ni-NTA affinity chromatography (Qiagen) in 50 mM Tris pH 8.0, 300 mM NaCl, 25 mM imidazole, 0.5 mM TCEP, 0.1 mM MnCl_2_; and eluted with 250 mM imidazole followed by size-exclusion chromatography on Superdex 75 increase 10/300 column (Cytiva) in 50 mM Tris pH 8.0, 150 mM NaCl, 0.5 mM TCEP, and 0.1 mM MnCl_2_. λPP activity was verified using a colorimetric pNPP assay, and aliquots were frozen at –80°C in 50% glycerol.

### In-vitro phosphorylation assays

In-vitro Ago2 phosphorylation assays were performed as established previously^6^. All RNAs were purchased from Dharmacon. In brief, 1 μM of the assembeled HsAgo2:guide:target complex was incubated in kinase reaction buffer (25 mM Tris pH 7.4, 10 mM MgCl_2_, 2.5 mM DTT, 0.5 mM Na_3_VO_4_) together with a total of 66 μM ATP mix containing 0.6 nM [γ-^32^P] ATP (Perkin-Elmer). The kinase reaction was started by adding 20 nM CK1α, incubated at 37°C for 90 mins, and quenched with stop buffer (25 mM EDTA and 25 mM unlabeled ATP). The aliquots were spotted on phosphocellulose paper, washed three times in 75 mM phosphoric acid and once in acetone (5 min each), air-dried, and measured using liquid scintillation counting. Moles phosphorylation per mole Ago2 were calculated and presented. (n ≥ 3; for all the experiments).

### Target binding assays

Equilibrium binding assays were performed to analyze RNA binding as previously described^6^ with minor modifications. A double-membrane filter was assembled with a top nitrocellulose membrane (to capture protein-RNA complex) and bottom nylon membrane (to capture free RNA) using the Bio-dot microfiltration system (Bio-Rad). RISC complexes were assembled by mixing Ago2 and guide (miR200 or miR200 variants) in 1:1.1 molar ratios and incubation at RT for 30 min in binding buffer (10 mM Tris pH 8.0, 100 mM NaCl, 5 mM DTT). Then 0.015 nM of the 5’ ^32^P labelled targets were mixed with different RISC concentrations (2-fold serial dilutions from 100 nM to 0.003 nM) in 100 µl reaction in binding buffer and incubated for 1 hr at RT before separating the RISC-bound and free-RNA fractions by slot-blot. The membranes were washed with 100 µl of binding buffer before drying and analyzing by phosphor imaging (Typhoon FLA 7000, Cytiva). Blot quantifications were done using ImageJ and analyzed using GraphPad Prism9. n>2 for all assays.

### Guide binding assay

0.015 nM of 5’ ^32^P labelled guide was incubated with different concentrations of Ago2 (or Ago2 mutants) (2-fold serial dilutions from 100 nM to 0.78 nM) in a 100 µl reaction in binding buffer (10 mM Tris pH 8.0, 100 mM NaCl, 5 mM DTT), and incubated for 1 hr at RT, before applying to the microfiltration system. The blots were processed and analyzed as described above.

### Slicing assays

Target slicing assays were carried out as described previously^27^ with minor modifications. In brief, RISC core was assembled by mixing 1 μM HsAgo2 and miR200b guide in 1:1 molar ratio and incubating at RT for 30 min in slicing buffer (10 mM Tris pH 8.0, 100 mM KCl, 10 mM DTT, 2 mM MgCl_2_, 10% glycerol). RISC (100 nM) was pre-incubated at 37°C for 5 min in 40 μl slicing buffer, and the reaction was started by adding 0.5 nM of 5’ ^32^P-labelled fully-complementary target. 4 μl reaction volume was collected at 0, 1, 2, 5, 10, 20, and 40 min time points and directly quenched with 4 μl stop buffer (80 % formamide, 0.1 % xylene cyanol, 0.1 % bromophenol blue, 2 mM EDTA, 1.5 M urea). Samples were heated to 95 °C for 5 min and analyzed on denaturing urea-PAGE. Gels were exposed overnight to phosphor screens and autoradiographed using Typhoon FLA 7000 (Cytiva). Quantification of substrate and product bands was done by ImageJ and data processed using GraphPad Prism-9 (n=3).

### Analytical size-exclusion chromatography

Analytical size-exclusion chromatography (aSEC) experiments were performed using the Superdex 200 increase 3.2/300 column (Cytiva) in aSEC buffer (25 mM Tris pH 8.0, 150 mM NaCl, 5 mM DTT, 1 mM MgCl_2_, and 5% glycerol). A RISC-target complex (RISC^ZT30^) was assembled by sequentially incubating 7.5 μM miR200b and 7.5 μM ZT1 target RNAs with 6.5 μM Ago2 for 5 min at RT in aSEC buffer. Next, 13 μM of CK1α (WT or mutant) were incubated with RISC^ZT30^ complex in the presence of 165 μM AMPPNP for 25 min at RT before injection into the SEC column. The resulting fractions corresponding to the complex peak were analyzed on 4-20% SDS-PAGE.

### Cryo-EM sample preparation

For cryo-EM grid preparation, 5 μM of purified HsAgo2 (in 25 mM HEPES pH 7.4, 100 mM NaCl, and 5 mM DTT) was incubated with miR200b guide in 1:1.2 molar ratio at room temperature (RT) for 30 min, followed by addition of ZT1 target RNA (1:1.2:1.2 molar ratio) for 30 min at RT. The assembled Ago2:guide:target complex was further incubated with CK1α (1:1.2:1.2:2 molar ratio) together with 100 μM AMPPNP and 2 mM MgCl_2_ for 1 hr. Since we did not observe complex particles on the micrographs likely due to their disintegration at the air-water interface during sample preparation, we employed mild crosslinking of the complex, a common approach to stabilize protein complexes for cryo-EM. We screened several crosslinkers, including DSG, BS3, DSS, glutaraldehyde, and formaldehyde and chose 0.25 mM of No-weigh format disuccinimidyl suberate (DSS) (Thermo Scientific Pierce) for 90 min at RT for our crosslinking protocol. The crosslinking reaction was quenched with 50 mM Tris pH 7.0, the sample concentrated and injected into the Superdex200 increase 10/300 (Cytiva) pre-equilibrated in 25 mM HEPES pH 7.4, 100 mM NaCl, and 5 mM DTT. The peak fractions containing the crosslinked complex were pooled, concentrated, and 0.05% w/v β-OG (Octyl β-D-glucopyranoside) (ThermoFisher Scientific) detergent was added before applying 4 μl sample onto glow-discharged cryo-EM grids, incubated for 10 sec at 20°C and 95% humidity, blotted for 3.1 s, and plunged into liquid ethane using an EM GP2 automatic plunge freezer (Leica). The UltrAuFoil R 0.6/1 Au 300 mesh grids were used for both datasets for Ago2-miR200-ZT30-CK1α and Quantifoil R 0.6/1 Cu 300 mesh grids for Ago2-miR200-ZT14-CK1α and Ago2-miR200-ZT13-CkK1α datasets.

### Cryo-EM data acquisition

Cryo-EM data were collected on a Titan Krios transmission electron microscope (ThermoFisher Scientific) operating at 300 keV. EPU data collection software (v 2.10.0.5) (ThermoFisher Scientific) was used, and dose-fractionated movies were collected using a K3 direct electron detector (Gatan) operating in electron counting mode.

**Ago2-miR200-ZT30-CK1α** (dataset-1): 30-framed movies were collected with an exposure rate of 2.04 e^−^/Å^2^/frame, resulting in a cumulative exposure of 61.2 e^−^/Å^2^. A total of 4,328 micrographs were collected at 105,000x magnification (0.856 Å/pixel) and defocus range of 0.6 to 2.2 μm.

**Ago2-miR200-ZT30-CK1α** (dataset-2): 30-framed movies were collected with an exposure rate of 2.26 e^−^/Å^2^/frame, resulting in a cumulative exposure of 67.8 e^−^/Å^2^. A total of 5,027 micrographs were collected at 105,000x magnification (0.856 Å/pixel) and defocus range of 0.7 to 2.2 μm.

**Ago2-miR200-ZT14-CK1α**: 30-framed movies were collected with an exposure rate of 2.55 e^−^/Å^2^/frame, resulting in a cumulative exposure of 76.65 e^−^/Å^2^. A total of 8,606 micrographs were collected at 105,000x magnification (0.856 Å/pixel) and defocus range of 0.6 to 2.0 μm.

**Ago2-miR200-ZT13-CK1α**: 30-framed movies were collected with an exposure rate of 2.52 e^−^/Å^2^/frame, resulting in a cumulative exposure of 75.5 e^−^/Å^2^. A total of 6,501 micrographs were collected at 105,000x magnification (0.856 Å/pixel) and defocus range of 0.7 to 2.2 μm in CDS (correlated-double sampling) mode.

### Cryo-EM Image processing

WARP (v 1.0.9)^44^ was used for real-time image pre-processing (motion correction^45,46^, CTF estimation^47,48^, and particle picking^49^) for all datasets. Particle picking was performed with the BoxNet pretrained neural network bundle included in WARP that is implemented in TensorFlow. A particle diameter of 120 Å and threshold score of 0.4 yielded 820,948 and 1,126,019 particle coordinates for Ago2-miR200-ZT30-CK1α dataset-1 and set-2 respectively, 1,338,929 particle coordinates for Ago2-miR200-ZT14-CK1α dataset, and 817,861 particle coordinates for Ago2-miR200-ZT13-CK1α dataset could be extracted respectively.

All subsequent processing steps were carried out in cryoSPARC v3.258^50^. For all structures, extracted particles were 2D classified, and a subset of those were used for ab-initio 3D reconstruction after manually inspecting each 2D class. We further separated particles in upto 11 ab-initio classes, and resulting 3D maps were heterogeneously refined against the whole particle set.

#### Ago2-miR200-ZT30-CK1α

259,567 and 199,374 particles (from dataset-1) and 330,628 and 250,772 particles (from dataset-2) could be separated into CK1α-bound and CK1α-free classes, respectively. The CK1α-bound particles from both datasets were combined (total 590,195 particles) and subsequently separated via a heterogeneous refinement against 4 junk ab-initio classes. Filtered 487,577 particles were refined homogeneously and then via a non-uniform refinement cycle to generate a map for the RISC^ZT30^-CK1a structure with global GSFSC resolution estimated up to 2.73 Å. The particles for RISC^ZT30^ classes from both datasets were also combined and processed similarly to the CK1α-bound particle sets, which yielded a cryo-EM map for RISC^ZT30^ with global GSFSC resolution of 2.8 Å estimated from 375,289 particles.

#### Ago2-miR200-ZT14-CK1α

336,017 and 436,695 particles could be segregated into CK1α-bound and CK1α-free classes. A subsequent round of heterogeneous refinement of these maps against 3 junk ab-initio classes separated 314,744 and 418,084 particles in CK1α-bound and CK1α-free classes. Next, one round of homogeneous and non-uniform refinement for the two particle classes was performed, which yielded cryo-EM maps for RISC^ZT14^-CK1α and RISC^ZT14^ with overall GSFSC resolutions of 3.7 Å and 3.4 Å, respectively.

#### Ago2-miR200-ZT13-CK1α

After particle sorting, two CK1α-free forms were predominantly observed in this dataset. A total of 239,359 and 167,512 particles were separated in two major CK1α-free classes. Careful inspection revealed that another class with 189,571 particles, showing RISC density also contains weak density for CK1α as well, processed as RISC^ZT13^-CK1α complex. An iterative round of heterogeneous refinement filtered out 221,592, 152,408, and 172,197 particles in three different maps. Subsequent homogeneous and a non-uniform refinement yielded a 3.4 Å map for RISC^ZT13^ closed, 3.7 Å map for RISC^ZT13^-open and 3.7 Å map for RISC^ZT13^-CK1α states, according to the GSFSC estimates. A local refinement masking CK1α was performed to improve the map density for the kinase in RISC^ZT13^-CK1α and exhibited only minor improvements.

We further performed local sharpening and de-noising of cryo-EM maps using non-linear post-processing with Deep cryo-EM Map Enhancer^51^. As previously reported^8^, these DeepEMhanc’ed maps had improved cryo-EM density, especially for the RNA, and were used to assist model building, while original maps were used for structure model refinements. To further examine RISC-CK1α states, we performed focused refinement^52^ with a 3D mask around the CK1α, but that did not improve the CK1α map quality. 3D variability analyses^53^ for different RISC datasets were performed using cryoSPARC to reveal dynamic movements of Ago2 domains across our different structures.

### Model-building, refinement, and validation

The crystal structures of the HsAgo2/miR122 complex excluding the target^20^ (PDB 6N4O) and CK1α^25^ (PDB 6GZD) were initially used for rigid body fitting in the cryo-EM map using ChimeraX^54^. The guide sequence was manually replaced to match the miR200b sequence, while the target RNA was built de-novo in the cryo-EM map. The RISC^ZT30^-CK1α structure was further used as a template for the other structures reported here. All atomic model building was done in Coot (v 0.9.4)^55^, and refinements were performed in PHENIX (v1.20.1-4487-000)^56^. Secondary structure restraints for protein and RNA were used throughout the refinement process. The DeepEMhance’d maps were also utilized for visualizing and building the structure models, while we used the unsharpened cryo-EM map for refinements. Structure validation was done using the MolProbity server^57^. The structure figures were generated by ChimeraX and PyMOL molecular graphics system (Version 2.5.5, Schrödinger, LLC, Heidelberg, D). Data collection and model statistics are summarized in Table S1.

## Acknowledgments

We thank members of Joshua-Tor laboratory for helpful discussions. We thank Dennis Thomas, the Director of the CSHL Cryo-EM facility for support with Cryo-EM data collection, and the CSHL Mass Spectrometry shared resource, supported by the CSHL Cancer Center Support Grant 5P30CA045508. We thank Steven F. Dowdy lab for discussions and support with chemical modifications of some of the RNAs. L.B. is supported by CSHL School of Biological Sciences. L.J. is an Investigator of the Howard Hughes Medical Institute.

## Author contributions

A.G. and L.J conceived the study. A.G. designed the constructs. A.G., L.B. and B.B. purified the proteins. A.G., L.B., A.A and B.B. performed the biochemical assays. A.G. and B.B. optimized the cryo-EM sample preparation conditions, and A.G. prepared the cryo-EM grids, collected the cryo-EM data and solved the structures. A.G. and L.J. analyzed the data and wrote the manuscript, and all authors edited the text.

## Data and materials availability

The generated cryo-EM maps and PDB codes associated with different structures have been deposited in the EMDB and PDB databases, with the details provided in **Supplementary table S1**. Further requests for resources and reagents generated in this study should be directed to Leemor Joshua-Tor.

