## Supplementary Material for "Structural dynamics between Argonaute-2 and CK1α promote target RNA release in microRNA-mediated silencing"

### Supplementary Figure 1-

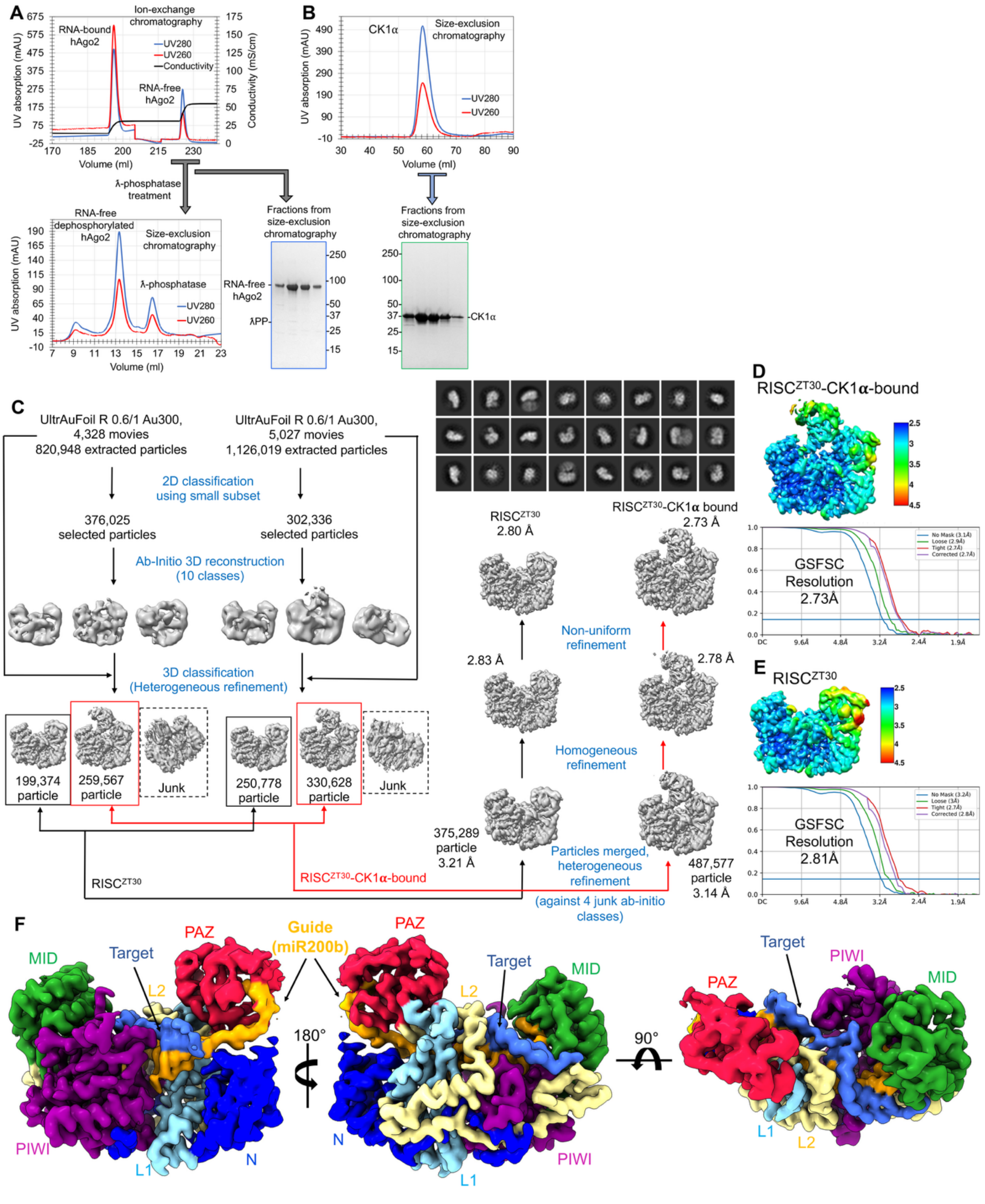

**Supplementary Figure 1-** Structural analysis of $RISC^{ZT30}$ in complex with CK1α kinase. (A) Purification of dephosphorylated RNA-free human Ago2 (HsAgo2). The ion exchange chromatogram showing separation of the RNA-bound Ago2 fraction from the RNA-free Ago2 fraction. The SEC chromatogram shows separation of the phosphatase-treated RNA-free Ago2 from the phosphatase. SDS-PAGE showing the purified Ago2 protein used in the study. (B) SEC chromatogram for the CK1α showing a homogeneous protein. The peak fractions analyzed on the SDS-PAGE showing pure protein used in the study. (C) CryoEM data processing workflow for the $RISC^{ZT30}$-CK1α complex. Processed particles from two different datasets were merged to obtain the $RISC^{ZT30}$ and $RISC^{ZT30}$-CK1α structures. The observed 2D class averages for some of the classes used in data processing are also shown. The local resolution estimates and gold-standard FSC (GSFSC) resolution estimate for (D) $RISC^{ZT30}$-CK1α and (E) $RISC^{ZT30}$ maps. (F) CryoEM map for the $RISC^{ZT30}$ in different orientations. Different domains in Ago2, guide, and target are colored and marked.

**Supplementary Figure 2-**

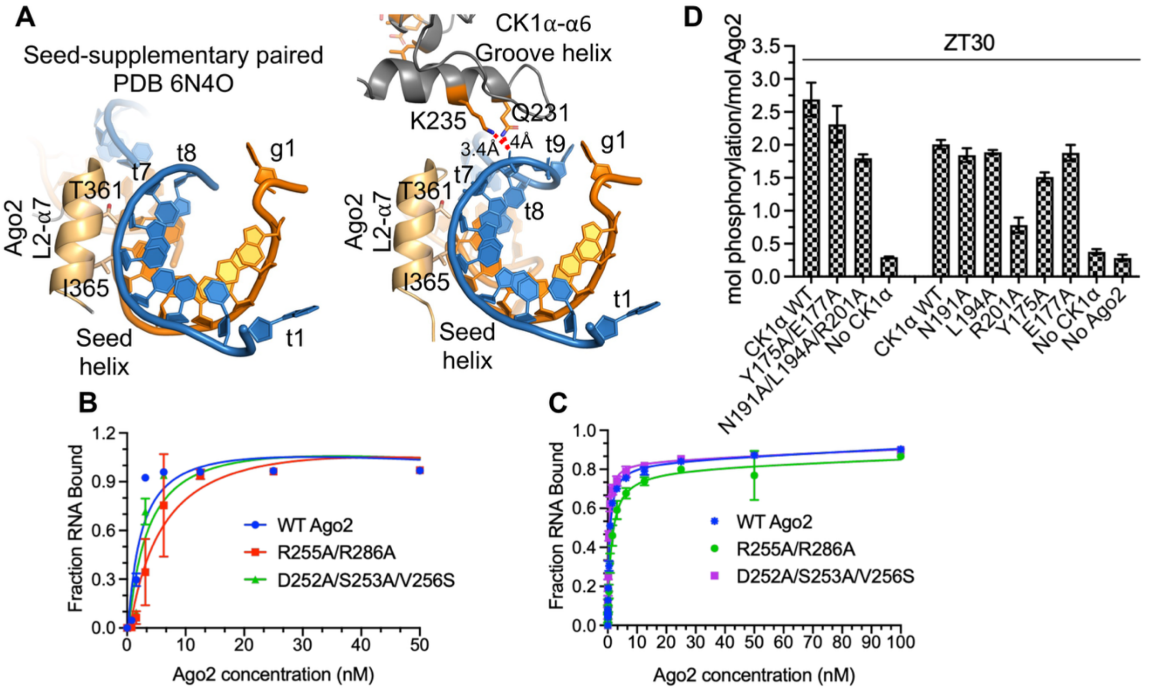


| Ago2 | Kd (nM) | Lower limit (95% CI) | Upper limit (95% CI) |
|---|---|---|---|
| WT | 2.2 | 1.4 | 3.7 |
| R255A/R286A | 6.0 | 3.3 | 11.5 |
| D252A/S253A/V256S | 3.3 | 2.0 | 5.7 |

| Ago2 | Kd (nM) | Lower limit (95% CI) | Upper limit (95% CI) |
|---|---|---|---|
| WT | 0.63 | 0.6 | 0.7 |
| R255A/R286A | 1.3 | 1.0 | 1.6 |
| D252A/S253A/V256S | 0.4 | 0.3 | 0.4 |

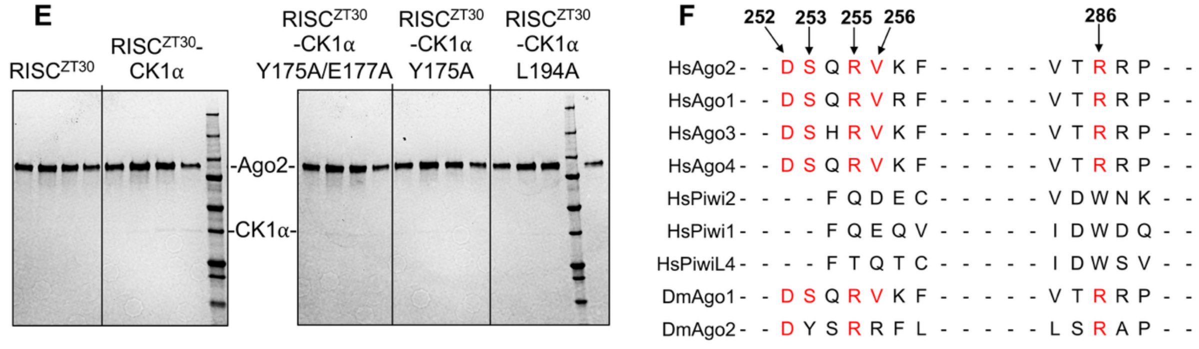


**Supplementary Figure 2-** CK1α interactions with RISC$^{ZT30}$. (A) Side-by-side view of the seed-helix supporting interactions in the Ago2 crystal structure with seed+supplementary region paired (PDB 6N4O)[1] and the RISC$^{ZT30}$-CK1α structure. CK1α groove-helix (α6) K235, Q231 establishes direct interaction with the guide-target seed-helix. The g6-g7 backbone kink disappears in RISC$^{ZT30}$. (B) miR200 guide and (C) ZT30 target binding assays for Ago2 PAZ mutants. The estimated RNA binding affinities (Kd) with 95% CI limits are also shown (n≥3). (D) *In vitro* Ago2 phosphorylation assay with ZT30 (checkered pattern) target RNA using different CK1α mutants at the CK1α-PAZ interface. Point mutations in R201 or in cluster mutants significantly reduced Ago2 phosphorylation. (E) SDS-PAGE gels showing the peak fractions for different aSEC experiments with $RISC^{ZT30}$ and CK1α mutants (F) Sequence alignment of PAZ domain residues that interact with CK1α in different Ago and Piwi proteins. These residues are completely conserved (red) in all human (Hs) Agos, but not in Piwi proteins or the slicing-specific fly (Dm)Ago2. Residue numbers correspond to HsAgo2.

#### Supplementary Figure 3-

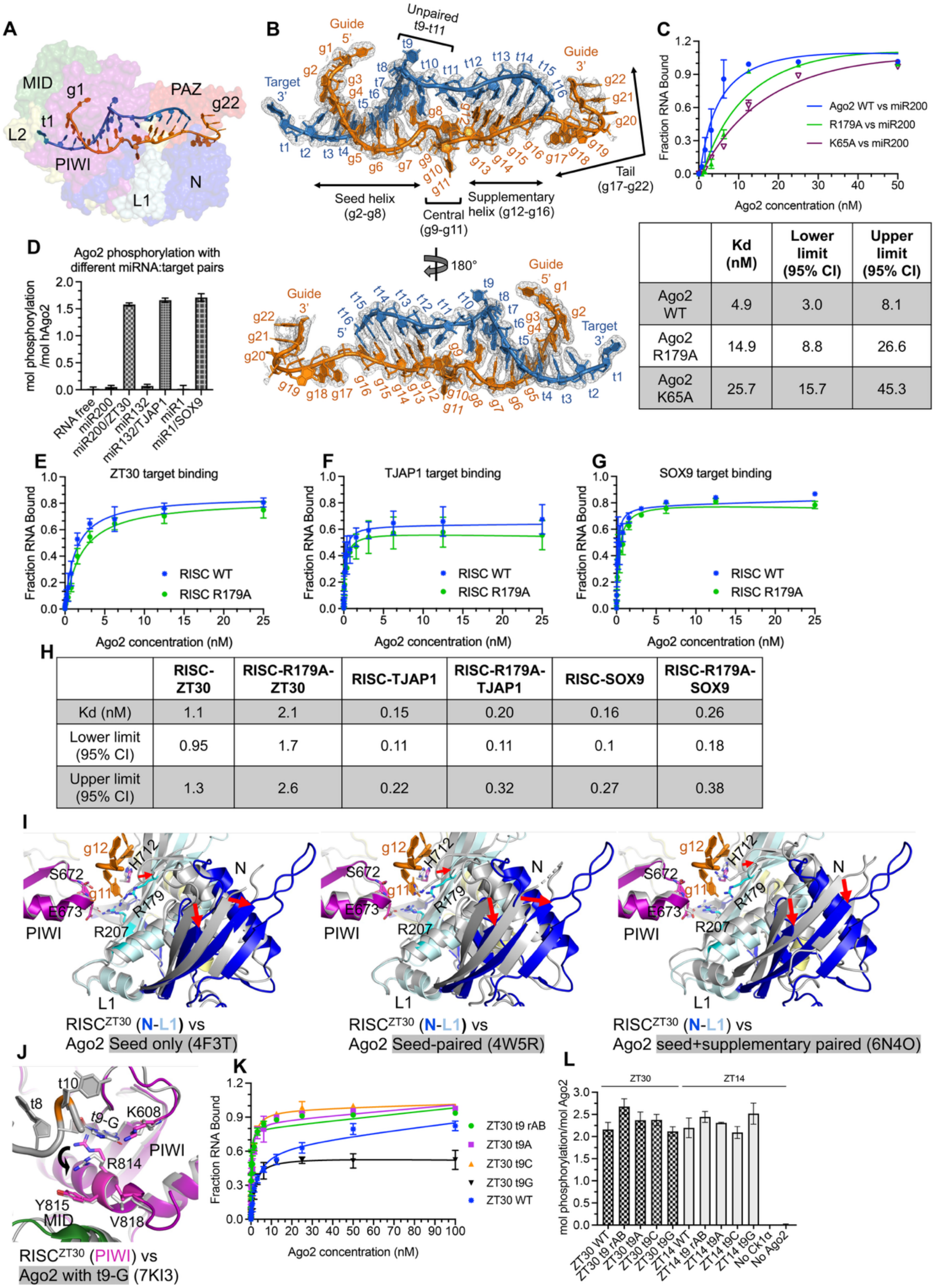


| | Kd (nM) | Lower limit (95% CI) | Upper limit (95% CI) |
|---|---|---|---|
| Ago2 WT | 4.9 | 3.0 | 8.1 |
| Ago2 R179A | 14.9 | 8.8 | 26.6 |
| Ago2 K65A | 25.7 | 15.7 | 45.3 |

| | RISC-ZT30 | RISC-R179A-ZT30 | RISC-TJAP1 | RISC-R179A-TJAP1 | RISC-SOX9 | RISC-R179A-SOX9 |
|---|---|---|---|---|---|---|
| Kd (nM) | 1.1 | 2.1 | 0.15 | 0.20 | 0.16 | 0.26 |
| Lower limit (95% CI) | 0.95 | 1.7 | 0.11 | 0.11 | 0.1 | 0.18 |
| Upper limit (95% CI) | 1.3 | 2.6 | 0.22 | 0.32 | 0.27 | 0.38 |

**Supplementary Figure 3-** Structural features of guide-target binding in RISC. (A) RISC$^{ZT30}$ structure showing the complete guide-target duplex. (B) The cryoEM map density for the duplex in two different orientations. (C) miR200 guide binding assay for Ago2 with the g11-stacking R179 and g16-stacking K65 mutants. The estimated binding affinities (Kd) with 95% CI limits are also shown (n≥3). (D) *In vitro* Ago2 phosphorylation assay testing a few different guide-target pairs. Guide binding alone shows no Ago2 phosphorylation, while guide-target binding triggers Ago2 phosphorylation (n≥3). Binding assays for (E) ZT30, (F) TJAP1 and (G) SOX9 target RNAs with RISC WT or R179A mutation (n=3 for ZT30 and TJAP1, n=2 for SOX9) (H) Estimated binding affinities (Kd) with 95% CI limits. (I) A comparative structural analysis showing superimposed g11 (gold) in N-L1 from RISC$^{ZT30}$ (colored cartoon) with previous Ago2 structures (grey cartoon) containing seed-only (PDB 4F3T)[2], seed-paired (PDB 4W5R)[3], or seed+supplementary paired (PDB 6N4O)[1]. g11 could fit in the same site in other Ago2 structures with appropriate movements in N-L1 domains (highlighted with red arrows). The g11 site is often occupied by R179, and L1 repositioning would open the pocket for g11 binding. (J) A superposition of the RISC$^{ZT30}$ (colored) and a previous Ago2 structure containing a G at the t9 position (grey) (PDB 7KI3)[4]. PIWI R814 would have to reposition in order to fit a t9-G nucleobase (black arrow). Other t9 pocket residues don't move. (K) ZT30 target binding in RISC using t9 RNA variants as indicated. Compared to other ZT30 targets, t9-G RNA shows impaired binding with a significant reduction in the target-bound species at a saturating RISC concentration (100 nM) in filter binding assays (n=3). (L) *In vitro* phosphorylation assay of Ago2 with t9 variants in ZT30 (checkered pattern) and ZT14 (grey) RNAs. Substitution of t9 did not affect Ago2 phosphorylation levels.

**Supplementary Figure 4-**

A

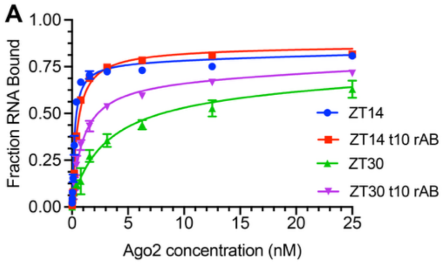


B

| RISC | Kd (nM) | Lower limit (95% CI) | Upper limit (95% CI) |
|---|---|---|---|
| ZT30 | 3.0 | 2.2 | 4.2 |
| ZT30 t10 rAB | 0.95 | 0.71 | 1.25 |
| ZT14 | 0.20 | 0.16 | 0.25 |
| ZT14 t10 rAB | 0.44 | 0.37 | 0.52 |

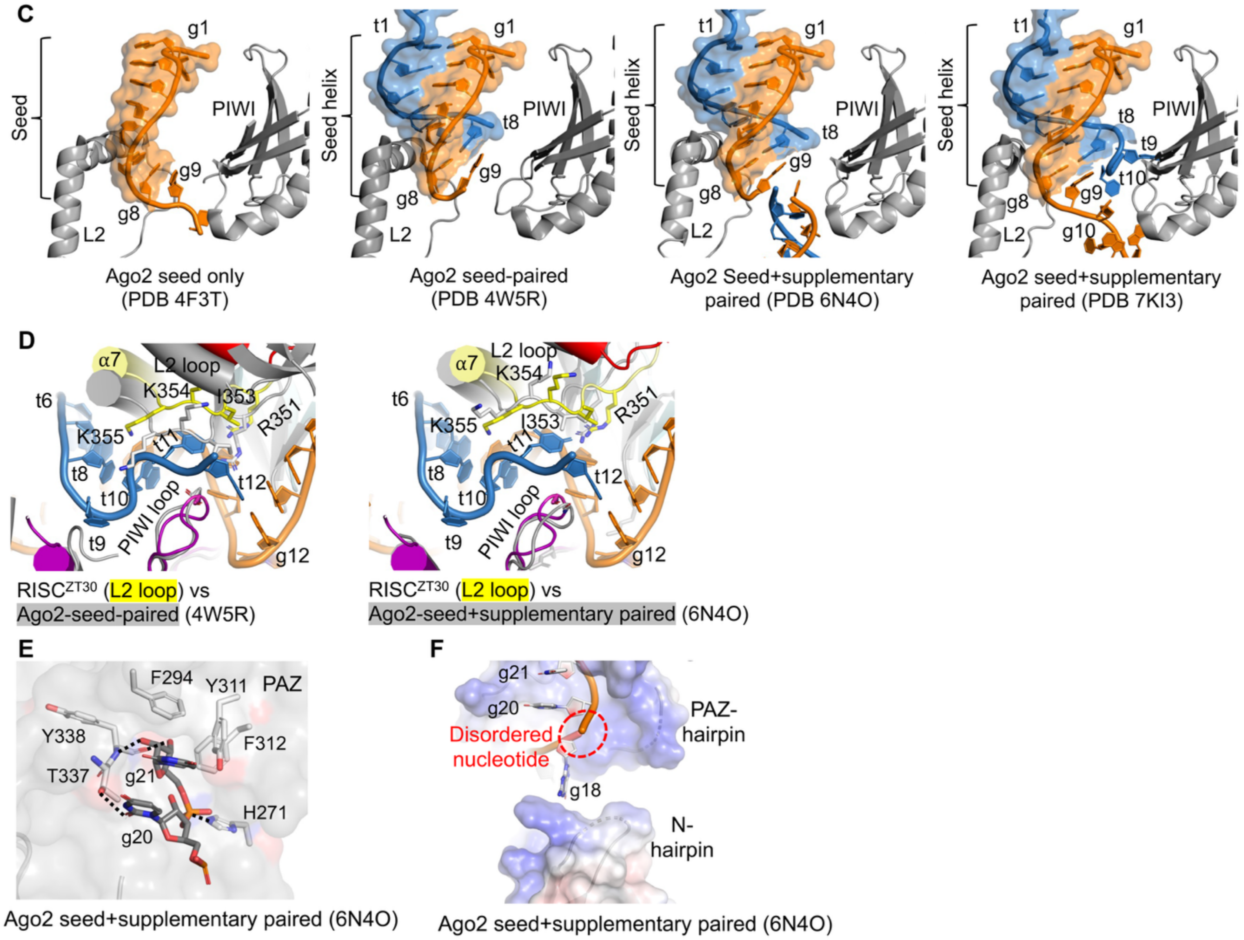


**Supplementary Figure 4-** Features in RISC-target complex assembly. (A) RISC-target binding assays of t10 variants in ZT30 and ZT14 target RNA. (B) Calculated binding affinities (Kd) with 95% CI limits (n=>2). RISC binding with a t10 abasic nucleotide (rAB) does not change binding significantly. (C) Analysis highlighting the g9 positioning in different Ago2 structures containing seed only (PDB 4F3T)[2], seed-paired (PDB 4W5R)[3], seed-supplementary paired (PDB 6N4O and 7KI3)[1,4]. The g9 nucleobase stacks under g8 or the g8-t8 base-pair supporting the seed or seed-helix region. (D) Superposition of the target RNA (blue) from the RISC$^{ZT30}$ structure on the seed-paired (PDB 4W5R) and seed-supplementary paired (PDB 6N4O) showing that t11 would clash with the L2 loop (R351-K355 region) in those previous Ago2 structures. (E) The 3' tail interaction with PAZ in Ago2 structure (PDB 6N4O). The g20-g21 are tightly bound in the PAZ 3' pocket. The direct H-bonds are highlighted with black dotted lines. (F) The open tail-gate region in the previous Ago2 structure containing seed-supplementary paired region (PDB 6N4O). g19 in this structure remains disordered.

**Supplementary Figure 5-**

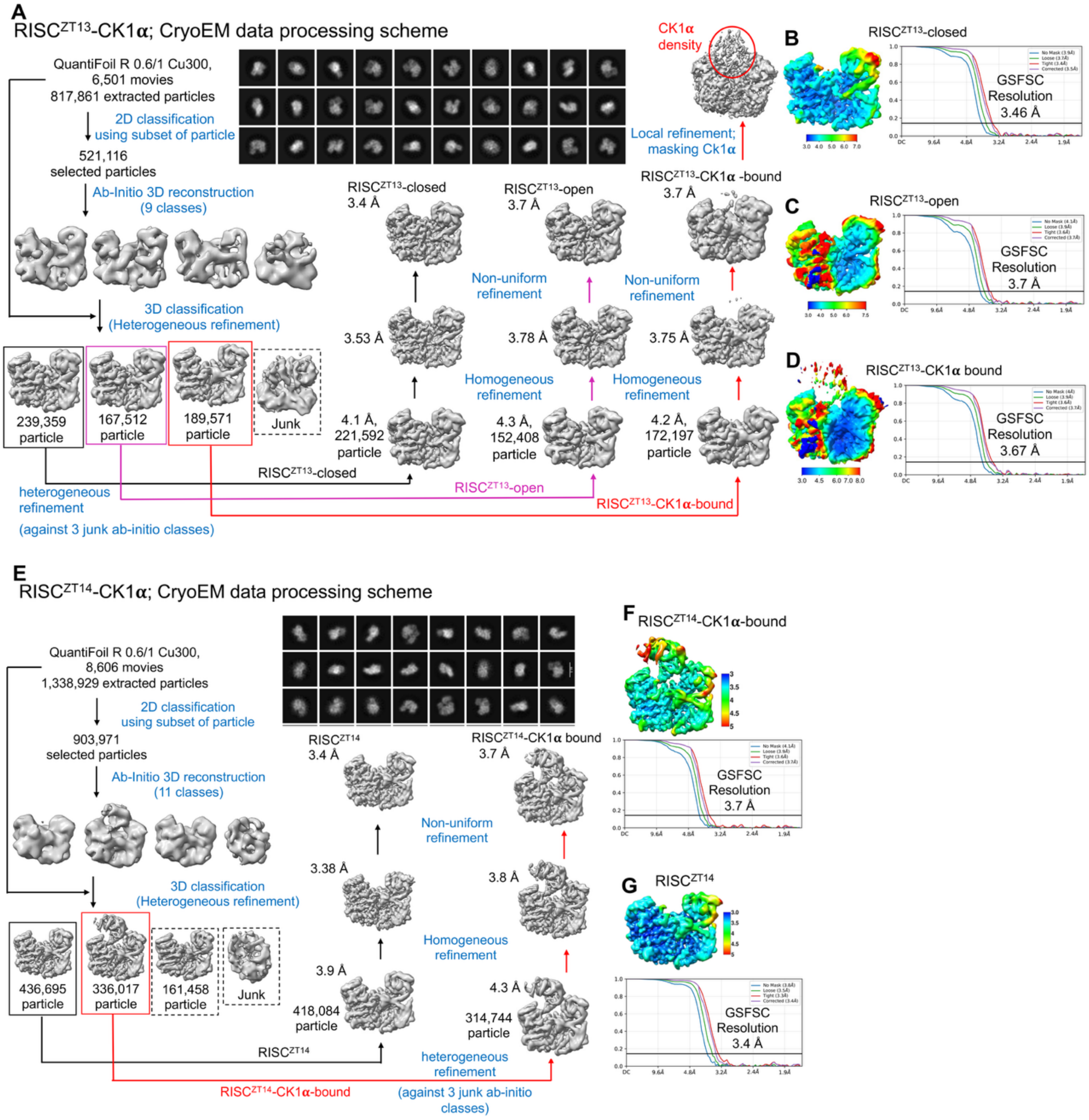


**Supplementary Figure 5-** Structural analysis of RISC complexes with shorter targets and CK1α kinase. (A) CryoEM data processing workflow for the RISC$^{ZT13}$-CK1α complex. Pre-processed cryoEM particles were separated into RISC$^{ZT13}$-closed, RISC$^{ZT13}$-open, and RISC$^{ZT13}$-CK1α states. The observed 2D class averages for some of the classes used in data processing are shown. Local resolution estimates and gold-standard FSC (GSFSC) resolution estimates for (B) RISC$^{ZT13}$-closed (C) RISC$^{ZT13}$-open and RISC$^{ZT13}$-

CK1α maps. (E) CryoEM data processing workflow for the RISC$^{ZT14}$-Ck1α complex, with 2D average class representatives. The local resolution estimates and gold-standard FSC (GSFSC) resolution estimates for (F) RISC$^{ZT14}$-Ck1α and (G) RISC$^{ZT14}$ EM maps.

#### Supplementary Figure 6-

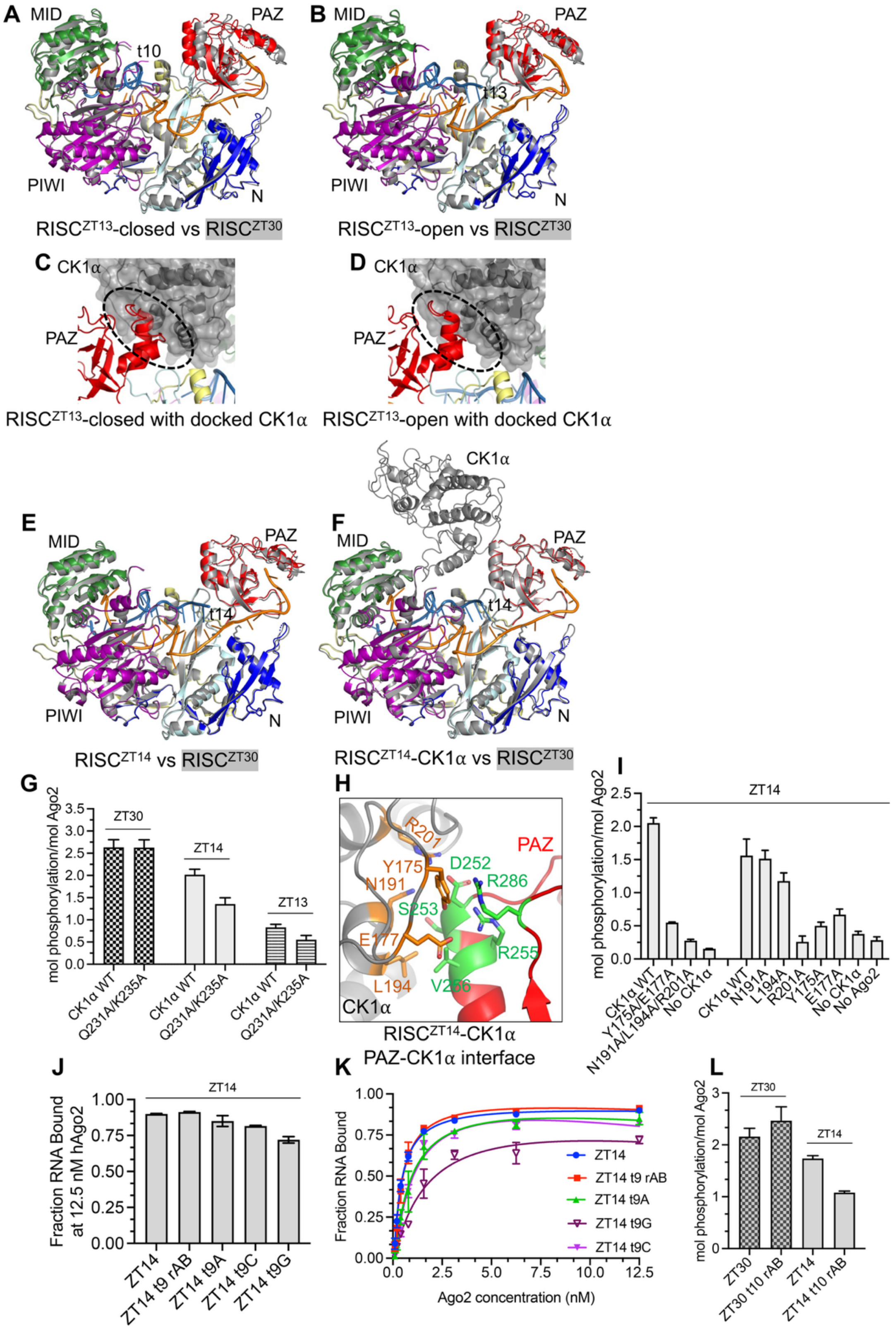

**Supplementary Figure 6**- RISC dynamics during CK1α docking. Structural superposition of the (A) RISC$^{ZT13}$- closed and (B) RISC$^{ZT13}$-open structures with the RISC$^{ZT30}$ (grey) structure. The PAZ domain leans toward the MID domain in the RISC$^{ZT13}$-closed structure and widens the RNA binding channel by swinging away from the MID domain. The complete ZT13 target (sky blue) is visible in the open structure, while the RNA in the closed structure is visible only up to the t10 nucleotide. (C) Superimposition of CK1α on the RISC$^{ZT13}$-closed structure showing clashes at the interface (black dotted circle). (D) Superimposition of CK1α on RISC$^{ZT13}$-open structure does not show clashes with the PAZ domain (black dotted circle). Superimpositions of (E) RISC$^{ZT14}$ and (F) RISC$^{ZT14}$-CK1α structures with RISC$^{ZT30}$ (grey). Both RISC$^{ZT14}$ structures superimpose well on RISC$^{ZT30}$ without any clashes between CK1α and PAZ. The complete ZT14 target (sky blue) and miR200 guide (yellow) are visible in both structures. (G) *In vitro* Ago2 phosphorylation assays with CK1α containing groove-helix mutations with different target RNA lengths: ZT30, ZT14, and ZT13. Ago2 phosphorylation is significantly reduced with the ZT14 target but not with other targets. (H) A zoomed-in view of the CK1α-PAZ interface in RISC$^{ZT14}$-CK1α. All the observed interactions are conserved in the RISC$^{ZT30}$-CK1α. (I) *In vitro* Ago2 phosphorylation assays using ZT14 (grey bar) target RNA with different CK1α mutations at the CK1α-PAZ interface. Different point mutations show an almost complete loss of Ago2 phosphorylation. (J) The calculated target-bound species at a saturating RISC concentration (12.5 nM) and the (K) traces for the binding assays for different ZT14 t9 RNA variants from filter binding assays (n=3). The t9-G substitution shows only a modest reduction in RISC binding in ZT14 target RNA. (L) *In vitro* Ago2 phosphorylation showing the effect of an abasic nucleotide at t10 (rAB) in the ZT14 and ZT30 targets. t10-rAB ZT14 shows reduced phosphorylation.

**Supplementary Fig 7**-

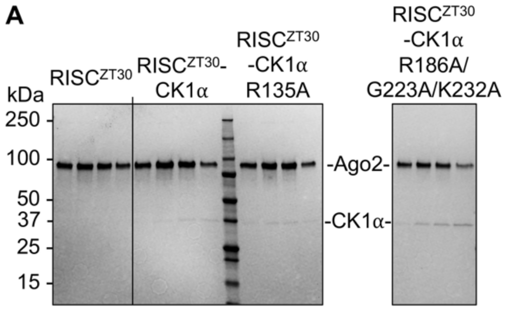


**Supplementary Figure 7**- CK1α ABS and RISC mutants. (A) SDS-PAGE gels analyzing peak fractions for different aSEC runs testing CK1α ABS mutant proteins and $RISC^{ZT30}$.

Supplementary movie 1- Movie showing the 3D variability observed for the $RISC^{ZT30}$-CK1α cryo-EM map in one orientation.

Supplementary movie 2- Movie showing the 3D variability observed for the $RISC^{ZT30}$-CK1α cryo-EM map in a second orientation.

Supplementary movie 3- Movie showing the 3D variability observed for the $RISC^{ZT14}$-CK1α cryo-EM map in one orientation.

Supplementary movie 4- Movie showing the 3D variability observed for the $RISC^{ZT14}$-CK1α cryo-EM map in a second orientation.

Supplementary movie 5- Movie showing the 3D variability observed for the $RISC^{ZT14}$ cryo-EM map.

**Supp. Table-1**: Data collection and refinement statistics for hAgo2-guide-target-CK1α cryo-EM structures

| **Cryo-EM sample** | **RISC$^{ZT13}$-CK1α** | | **RISC$^{ZT14}$-CK1α** | | **RISC$^{ZT30}$-CK1α** | |
|---|---|---|---|---|---|---|
| **Data collection and processing** | | | | | | |
| Microscope | Titan Krios | | | | | |
| Voltage (kV) | 300 | | | | | |
| Detector | K3 | | | | | |
| Magnification | 105,000x | | | | | |
| Pixel size (Å) | 0.856 | | | | | |
| Data collection software | EPU | | | | | |
| Defocus range (µm) | 0.7 – 2.2 | | 0.6 – 2.0 | | 0.6 – 2.2 | 0.7 – 2.2 |
| Total exposure ($e^-/Å^2$) | 75.5 | | 76.65 | | 61.2 | 67.8 |
| Frames/exposure | 30 | | | | | |
| Exposure/frame ($e^-/Å^2$) | 2.52 | | 2.55 | | 2.04 | 2.26 |
| Micrographs collected | 6,501 | | 8,606 | | 4,328 | 5,027 |
| Total extracted particles | 817,861 | | 1,338,929 | | 820,948 | 1,126,019 |
| **Name** | **RISC$^{ZT13}$-closed** | **RISC$^{ZT13}$-open** | **RISC$^{ZT14}$** | **RISC$^{ZT14}$-CK1α** | **RISC$^{ZT30}$** | **RISC$^{ZT30}$-CK1α** |
| PDB/EMDB | 10JF/75215 | 10JG/75216 | 10JH/75217 | 10JI/75218 | 10JJ/75219 | 10JK/75220 |
| Final particles | 221,592 | 152,408 | 418,084 | 314,744 | 375,289 | 487,577 |
| Symmetry | C1 | | | | | |
| **Map resolution (unmasked/masked)** | | | | | | |
| GSFSC 0.143 | 3.9/3.5 | 4.1/3.7 | 3.8/3.4 | 4.1/3.7 | 3.2/2.8 | 3.1/2.8 |
| **Refinement and model validation** | | | | | | |
| **Model resolution (unmasked/masked)** | | | | | | |
| FSC 0.143 | 3.6/3.5 | 3.8/3.7 | 3.4/3.3 | 3.7/3.7 | 2.8/2.8 | 2.8/2.7 |
| FSC 0.5 | 4.1/3.9 | 4.3/4.2 | 3.7/3.5 | 4.1/4.0 | 3.1/3.0 | 3.1/3.0 |
| Map CC mask/volume | 0.75/0.75 | 0.69/0.69 | 0.78/0.78 | 0.68/0.69 | 0.85/0.84 | 0.78/0.78 |
| Protein residue | 814 | 823 | 823 | 1111 | 825 | 1120 |
| Nucleic acid | 32 | 35 | 36 | 36 | 38 | 38 |
| Water/Ligand | 0/0 | 0/0 | 0/0 | 0/0 | 0/0 | 0/1 |
| **B-factors (Å$^2$)** | | | | | | |
| Protein | 67.31 | 67.11 | 67.09 | 64.20 | 67.02 | 64.08 |
| Nucleotide | 68.15 | 69.65 | 72.02 | 72.02 | 73.28 | 73.28 |
| **RMS Deviations** | | | | | | |
| Bond length (Å) | 0.002 | 0.003 | 0.003 | 0.002 | 0.006 | 0.006 |
| Bond angle (°) | 0.585 | 0.593 | 0.536 | 0.579 | 0.573 | 0.642 |
| **Ramachandran statistics** | | | | | | |
| Favored (%) | 92.29 | 95.59 | 96.21 | 96.73 | 94.76 | 95.51 |
| Allowed (%) | 7.34 | 4.41 | 3.79 | 3.27 | 5.24 | 4.49 |
| Outlier (%) | 0.37 | 0.0 | 0.0 | 0.0 | 0.0 | 0.0 |
| Rotamer outliers (%) | 0.0 | 0.0 | 0.0 | 0.1 | 0.14 | 1.01 |
| CaBLAM outliers (%) | 4.28 | 2.59 | 1.85 | 1.92 | 2.33 | 1.71 |
| Clash score (all atoms) | 8.91 | 10.73 | 4.66 | 7.09 | 4.84 | 5.67 |
| MolProbity score | 1.95 | 1.86 | 1.46 | 1.60 | 1.60 | 1.62 |
| FSC is Fourier shell correlation, and RMSD is root-mean-square deviation. | | | | | | |

**Supplementary table 2**- The sequence of RNA substrates used in the study.

| S.no. | Name | Details | Sequence (5'-3') |
|---|---|---|---|
| 1 | Guide | miR200 (22nt length) | UAAUACUGCCUGGUAAUGAUGA |
| 2 | ZT30 | 30 nt ZT1 target | ACAUUAGCUGAUUUUUACCUAUCAGUAUUA |
| 3 | ZT14 | 14 nt ZT1 target | ACCUAUCAGUAUUA |
| 4 | ZT13 | 13 nt ZT1 target | CCUAUCAGUAUUA |
| 5 | ZT30 t9 rAB | 30 nt ZT1 target with rAB at t9 | ACAUUAGCUGAUUUUUACCUA(rAB)CAGUAUUA |
| 6 | ZT30 t9A | 30 nt ZT1 target with t9A substitution | ACAUUAGCUGAUUUUUACCUAACAGUAUUA |
| 7 | ZT30 t9C | 30 nt ZT1 target with t9C substitution | ACAUUAGCUGAUUUUUACCUACCAGUAUUA |
| 8 | ZT30 t9G | 30 nt ZT1 target with t9G substitution | ACAUUAGCUGAUUUUUACCUAGCAGUAUUA |
| 9 | ZT14 t9 rAB | 14 nt ZT1 target with rAB at t9 | ACC UA(rAB) CAG UAU UA |
| 10 | ZT14 t9A | 14 nt ZT1 target with t9A substitution | ACCUAACAGUAUUA |
| 11 | ZT14 t9C | 14 nt ZT1 target with t9C substitution | ACCUACCAGUAUUA |
| 12 | ZT14 t9G | 14 nt ZT1 target with t9G substitution | ACCUAGCAGUAUUA |
| 13 | ZT30 t10 rAB | 30 nt ZT1 target with rAB at t10 | ACAUUAGCUGAUUUUUACCU(rAB)UCAGUAUUA |
| 14 | ZT14 t10 rAB | 14 nt ZT1 target with rAB at t10 | ACC U(rAB)U CAG UAU UA-OH |
| 15 | ZTs | 30 nt ZT1 target with full complementarity to miR200 guide | ACAUUAGCUCAUCAUUACCAGGCAGUAUUA |
| 16 | g11 rAB | miR200 guide with rAB at g11 | UAAUACUGCC(rAB)GGUAAUGAUGA |
| 17 | g13/g14 rAB | miR200 guide with rAB at g13/g14 both | UAAUACUGCCUG(rAB)(rAB)AAUGAUGA |
| 18 | | | |

The red highlighted nucleotide positions are indicative of either mutations or substitution at that position in RNA sequence.

**Supplementary table 3**- List of the used reagents, generated plasmids, deposited PDB data and programs used in the study.

| **Reagent or Resource** | **Source** | **Identifier** |
|---|---|---|
| **Chemicals, peptides and recombinant proteins** | | |
| [γ-$^{32}$P] ATP | Revvity | Cat# BLU502Z250UC, BLU035C001MC |
| Strep-Tactin 4flow high-capacity resin | IBA Lifesciences | Cat# 2-1250-025 |
| SYBR Gold Nucleic acid gel stain | ThermoFisher Scientific | Cat# S11494 |
| DSS, No-weight format | ThermoFisher Scientific | Cat# A39267 |
| T4 Polynucleotide Kinase | NEB | Cat# M0201L |
| Proteinase K | NEB | Cat# P8107S |
| RNasIn RNase inhibitor | Promega | Cat# N2515 |
| Octyl β-D-glucopyranoside | ThermoFisher Scientific | Cat# BP585-1 |
| HyClone CCM3 media | Cytiva | Cat# SH30065.02 |
| d-Desthiobiotin | Millipore-Sigma | Cat# D1411-1G |
| HiTrap SP-HP Column | Cytiva | Cat# 17115201 |
| Mono S 10/100 GL Column | Cytiva | Cat# 17516901 |
| Superose 6 increase 3.2/300 column | Cytiva | Cat# 29091598 |
| Superose 6 increase 10/300 column | Cytiva | Cat# 29091596 |
| Superdex 200 increase 3.2/300 column | Cytiva | Cat#28990946 |
| Superdex 200 increase 10/300 column | Cytiva | Cat# 28990944 |
| Superdex 75 increase 10/300 column | Cytiva | Cat# 29148721 |
| **Deposited data** | | |

| | | |
|---|---|---|
| Coordinates for Ago2-miR200b-ZT13 (RISC$^{ZT13}$)-open structure | This work | PDB:10JG |
| Cryo-EM maps for Ago2-miR200b-ZT13 (RISC$^{ZT13}$)-open structure | This work | EMDB:75216 |
| Coordinates for Ago2-miR200b-ZT13 (RISC$^{ZT13}$)-closed structure | This work | PDB:10JF |
| Cryo-EM maps for Ago2-miR200b-ZT13 (RISC$^{ZT13}$)-closed structure | This work | EMDB:75215 |
| Coordinates for Ago2-miR200b-ZT14 (RISC$^{ZT14}$) structure | This work | PDB:10JH |
| Cryo-EM maps for Ago2-miR200b-ZT14 (RISC$^{ZT14}$) structure | This work | EMDB:75217 |
| Coordinates for Ago2-miR200b-ZT14 (RISC$^{ZT14}$) -CK1α structure | This work | PDB:10JI |
| Cryo-EM maps for Ago2-miR200b-ZT14 (RISC$^{ZT14}$) -CK1α structure | This work | EMDB:75218 |
| Coordinates for Ago2-miR200b-ZT30 (RISC$^{ZT30}$) structure | This work | PDB:10JJ |
| Cryo-EM maps for Ago2-miR200b-ZT30 (RISC$^{ZT30}$) structure | This work | EMDB:75219 |
| Coordinates for Ago2-miR200b-ZT30 (RISC$^{ZT30}$) -CK1α structure | This work | PDB:10JK |

| | | |
|---|---|---|
| Cryo-EM maps for Ago2-miR200b-ZT30 ($RISC^{ZT30}$) - CK1α structure | This work | EMDB:75220 |
| **Experimental models: Cell lines** | | |
| Sf9 insect cells | ThermoFisher Scientific | Cat# 11496015 |
| High Five insect cells | ThermoFisher Scientific | Cat# B85502 |
| *E.coli* DH5α cells (high efficiency) | ThermoFisher Scientific | Cat# 18258-012 |
| *E.coli* DH10Bac cells | ThermoFisher Scientific | Cat# 12033-015 |
| *E.coli* Rosetta 2 (DE3) cells | Novagen | Cat# 71400 |
| **Oligonucleotides** | | |
| RNA sequences. See Table S2. | N/A | N/A |
| **Recombinant DNA** | | |
| pFL-SST-HsAgo2 | Elkayam et al., 2012 | N/A |
| pFL-SST-HsAgo2(R255A/R286A) | This work | N/A |
| pFL-SST-HsAgo2(D252A/S253A/V256S) | This work | N/A |
| pFL-SST-HsAgo2(D358A) | This work | N/A |
| pFL-SST-HsAgo2(R255A) | This work | N/A |
| pFL-SST-HsAgo2(R286A) | This work | N/A |
| pFL-SST-HsAgo2(D252A) | This work | N/A |
| pFL-SST-HsAgo2(S253A) | This work | N/A |
| pFL-SST-HsAgo2(V256S) | This work | N/A |
| pFL-SST-HsAgo2(K65A) | This work | N/A |
| pFL-SST-HsAgo2(R179A) | This work | N/A |
| pFL-SST-HsAgo2(R179A/K65A) | This work | N/A |
| pFL-SST-CK1α | Bri et al., 2022 | N/A |
| pFL-SST-CK1α (Y175A/E177A) | This work | N/A |

| | | |
|---|---|---|
| pFL-SST-CK1α (N191A/L194A/R201A) | This work | N/A |
| pFL-SST-CK1α (Y175A) | This work | N/A |
| pFL-SST-CK1α (E177A) | This work | N/A |
| pFL-SST-CK1α (N191A) | This work | N/A |
| pFL-SST-CK1α (L194A) | This work | N/A |
| pFL-SST-CK1α (R201A) | This work | N/A |
| pFL-SST-CK1α (R185A) | This work | N/A |
| pFL-SST-CK1α (R185A/G223A/K232A) | This work | N/A |
| pFL-SST-CK1α (R134A) | This work | N/A |
| λPP-pET28a | Bri et al, 2022 | N/A |
| **Software and algorithms** | | |
| EPU software | ThermoFisher Scientific | |
| CryoSPARC v2 | Punjani et al., 2017 | https://cryosparc.com |
| WARP | Tegunov and Cramer, 2019 | http://www.warpem.com/warp/ |
| Coot | Emsley and Cowtan, 2004 | https://www2.mrc-lmb.cam.ac.uk/personal/pemsley/coot/ |
| PHENIX 1.20 | Adams et al., 2010 | http://www.phenix-online.org/ |
| ChimeraX | Pettersen et al., 2004 | https://www.cgl.ucsf.edu/chimerax/ |
| PyMol 2.5.5 | Schrödinger LLC | https://pymol.org/2/ |
| MolProbity | Chen et al., 2010. | http://molprobity.biochem.duke.edu/ |
| PRISM 9.5.1 software | GraphPad | https://www.graphpad.com/scientific-software/prism/ |
